# A Prime/Pull RR2/CXCL11 Therapeutic Vaccine that Bolsters the Number and Function of Dorsal Root Ganglia Tissue-Resident HSV-Specific CD8^+^ T_RM_ Cells Protects Latently Infected Guinea Pigs from Recurrent Genital Herpes

**DOI:** 10.1101/2022.07.22.501208

**Authors:** Nisha Dhanushkodi, Swayam Prakash, Ruchi Srivastava, Pierre-Gregoire A. Coulon, Hawa Vahed, Latifa Zayou, Afshana Quadiri, Hubert Schaefer, Lbachir BenMohamed

## Abstract

Reactivation of herpes simplex virus type 2 (HSV-2) from latently infected dorsal root ganglia (DRG) and subsequent virus shedding in the genital tract trigger recurrent genital herpes. Memory CD8^+^ T cells play a critical role in preventing HSV-2 reactivation from latently infected DRG, thus reducing recurrent genital lesions. The role of T-cell attracting chemokines in promoting CD8^+^ T cell protective immunity in recurrent genital herpes remains to be fully elucidated. In this study, we investigated whether and how the CXCL11/CXCR3 pathway affects the frequency and function of DRG-resident CD8^+^ T cells and the severity of recurrent genital herpes. Latently infected guinea pigs were primed with the HSV-1 RR2 protein, delivered intramuscularly with CpG/Alum adjuvants, and the induced T cells were “pulled” from the periphery into the latently infected DRG using T-cell attracting CXCL11 chemokine, delivered to DRG. In the guinea pigs that received the prime/pull vaccine, we detected a significant increase in both the number and function of tissue-resident IFN-γ^+^CD103^+^CD44^+^CXCR3^+^CD8^+^ T_RM_ cells that infiltrated healed sites of the vaginal mucosa (VM) and DRG tissues. This was associated with a significant decrease in virus shedding and a reduction in both the severity and frequency of recurrent genital herpes lesions. In contrast, in the guinea pigs that received the RR2 vaccine alone, we detected fewer functional CD8^+^ T_RM_ cells and no reduction in the severity of recurrent genital herpes. These findings highlight the role of the CXCL11/CXCR3 chemokine pathway in shaping tissue-resident CD8^+^ T_RM_ cell protective immunity against recurrent genital herpes.

**IMPORTANCE:** Recurrent genital herpes is a common sexually transmitted disease worldwide. Currently, no FDA-approved therapeutic vaccines are available. In the present study, we used HSV-2 latently infected guinea pig to investigate a novel therapeutic prime/pull vaccine strategy based on priming T cells systemically, with a recombinantly expressed herpes envelope and tegument protein RR2 and “pulling” primed T cells into the tissues of latently infected ganglia with the T-cell-attracting chemokine, CXCL11. We discovered that this RR2/CXCL11 prime/pull vaccine elicited a significant reduction in virus shedding and a decrease in both the severity and frequency of recurrent genital herpes sores. This protection correlated with increased numbers of functional tissue-resident IFN-γ^+^CD103^+^CD44^+^CXCR3^+^CD8^+^ T_RM_ cells that infiltrate healed sites of the VM tissues and DRG. Our findings shed light on the role of T_RM_ cells in protection against recurrent genital herpes and propose the prime/pull therapeutic vaccine as a new strategy against genital herpes.

**TWEET:** The present study presents a novel RR2/CXCL11 prime/pull therapeutic vaccine that elicited a significant reduction in virus shedding and a decrease in both the severity and frequency of recurrent genital herpes sores.

## INTRODUCTION

Herpes simplex virus type 2 (HSV-2) affects more women than men, with a staggering 315 million, 5-49 years old women, currently infected (1–3). After exposure of the vaginal mucosa (VM), HSV-2 replicates in the mucosal epithelial cells thereby causing acute genital herpetic lesions (3–8). Once the acute primary infection is cleared, the virus enters the nerve termini innervating peripheral vaginal tissues and is subsequently transported to the nucleus of the sensory neurons of the dorsal root ganglia (DRG) entering a lifelong “steady-state” latency (7, 8).

Tissue-resident memory CD8^+^ T cells appear crucial in reducing virus reactivation from latently infected DRG and virus replication in the VM mucosa. In the absence of a strong local CD8^+^ T response in DRG, the virus reactivates from latently infected sensory neurons and sheds back into the genital tract, where it causes severe recurrent ulcerative lesions. Roughly 80% of HSV-2- seropositive women are asymptomatic unaware of their infection as they never develop any apparent recurrent symptoms (2). In contrast, in symptomatic women, the latent infection is often interrupted by sporadic reactivation leading to recurrent genital lesions and painful blisters that can burst and form ulcers (7). Despite the availability of many intervention strategies, such as sexual behavior education, barrier methods, and anti-viral drug therapies (e.g., Acyclovir and derivatives), eliminating or at least reducing recurrent genital herpes continues to be a challenge (3, 9–12). Besides, current antiviral drugs can neither prevent *de novo* infection or reactivation nor clear the latent virus. Moreover, both symptomatic and asymptomatic women experience subclinical virus shedding and, hence, can transmit the virus, underscoring the need for an anti-viral therapeutic vaccine to prevent or reduce virus reactivation and/or its shedding in the genital tract. Currently, the medical opinion is highly focused on an effective anti-viral therapeutic vaccine that would constitute the best approach to protect from recurrent genital herpetic disease (5, 13). It is becoming increasingly clear that the acquired immune responses that develop following the initial virus exposure are not sufficient for protection against recurrent genital herpes in symptomatic women (14–16). This implies that a successful therapeutic vaccine must be able to boost an immune response that is stronger and/or different than the acquired immunity induced by the virus itself (17). Studies have demonstrated that whole live attenuated vaccines induced B- and T-cell protective immunity against acute genital HSV-2 challenges in animal models (18). However, the same level of protection has yet to be achieved safely in clinical trials (19, 20). Protein-based therapeutic vaccines have proven to be excellent vaccine candidates due to their safety, cost-effectiveness, and rapid preparation (21). Interestingly, over the last two decades, only a single subunit protein vaccine strategy, based on the HSV-2 glycoprotein D (gD), delivered with or without glycoprotein B (gB), has been continuously tested in clinical trials (20, 22). This subunit vaccine strategy proved unsuccessful in clinical trials despite inducing strong neutralizing antibodies (20). Previous studies have identified other antigenic tegument proteins by screening positive HSV-2 open-reading frames (ORFs) using antibodies and T cells from HSV-2 seropositive individuals (23). However, aside from three reports, first by our group in 2012 (24, 25) and later by Genocea Biosciences, Inc. in 2014 (26), a comparison of the repertoire of HSV-2 proteins, encoded by the 84^+^ open reading frames- (ORF-) in the 152kb genome of the HSV-2, recognized by antibodies and T cells from HSV-2 seropositive symptomatic versus asymptomatic individuals is largely unknown.

Over the last five years, our group has made significant progress in identifying candidate HSV- 2 antigens and characterized the phenotype and function of tissue-resident T cells associated with protection. We found that: (*i*) HSV-2 protein RR2 (Ribonucleotide Reductase 2) was frequently and highly recognized by antibodies and T cells from the naturally “protected” asymptomatic individuals (*ii*) RR2 protein was the main HSV-2 antigen recognized by tissue-resident memory CD8^+^ T_RM_ cells, reside within both the DRG and vaginal mucocutaneous tissues of protected guinea pigs.

Due to the ethical and practical limitations in obtaining fresh DRG- and vaginal mucosa from herpes patients, a routine characterization of both the phenotype and function of human tissues- derived T_RM_ cells remains a challenge (27–29). Thus, a reliable small animal model that would mimic recurrent genital herpes, as it occurs in humans, would help to speed up the characterization of tissue-resident HSV-specific CD8^+^ T_RM_ cells and compare the contribution of DRG- and VM-derived versus peripheral blood-derived circulating CD8^+^ T cells in protection against recurrent genital herpes. Guinea pigs have been the widely used animal model of choice for pre-clinical testing of therapeutic vaccine candidates against recurrent genital herpes (30–32). Unlike the mouse model, intravaginal HSV-2 infections in guinea pigs: (*i*) lead to latency followed by intermittent, spontaneous reactivation events resulting in recurrent virus shedding in the genital tract, as occurs in women; and (*ii*) this virus shedding can develop into clinical and pathological recurrent genital lesions in symptomatic guinea pigs, with self-limiting quantifiable vulvovaginitis, akin to those observed in symptomatic women (30–32).

In this study, we used the guinea pig model of recurrent herpes to test the safety, CD8^+^ T_RM_ cell immunogenicity, and protective efficacy of recombinantly expressed RR2 protein (encoded by the *UL40* gene). RR2 protein was rationally chosen for this study as being the most highly and selectively recognized HSV-2 protein by effector CD8^+^ T cells from naturally “protected” asymptomatic women who, despite being infected, never develop any recurrent herpetic disease. We demonstrated that therapeutic prime/pull vaccination with RR2 and CXCL11 boosted the numbers of functional tissue-resident IFN-γ^+^CRTAM^+^CD8^+^ T_RM_ cells, that were localized to healed sites of the vaginal mucocutaneous (VM) tissues. The latter leads to a significant reduction in virus shedding and a decrease in both the severity and frequency of recurrent genital herpes lesions. In this report, we discuss the role of tissue-resident T_RM_ cells in the protection against recurrent genital herpes in the guinea pig model using herpes subunit vaccine candidates in combination with T-cell attracting chemokines.

## MATERIALS AND METHODS

### Animals

Female guinea pigs (*Hartley strain*, Charles River Laboratories, San Diego, CA) weighing 275-350 g (5-6 weeks old) were housed at the University of California, Irvine vivarium. The Institutional Animal Care and Use Committee of the University of California, Irvine reviewed and approved the protocol for these studies (IACUC # AUP-19-111). The number of animals required per group for each experiment was calculated based on our prior experience of using animal models in spontaneous and induced recurrent herpes infection and disease. A group size of 6 had 80% power to detect a difference of two-fold or higher between experimental group means with a significance level of 0.05.

### Vaccine candidates

We selected recombinantly expressed RR2 protein from HSV-2 as an antigen that is highly and selectively recognized by CD8^+^ T cells from naturally “protected” asymptomatic individuals.

### Infection and prime/pull vaccination in Guinea Pigs

Throughout this study, we used the MS strain of HSV-2, generously gifted by Dr. David Bernstein (Cincinnati Children’s Hospital Medical Center, University of Cincinnati, OH). Guinea pigs (*n* = 18) were infected intravaginally with 5 x 10^5^ pfu of HSV-2 (strain MS). Once the acute infection was resolved, latently infected animals were randomly divided into 2 groups (*n* = 6). Latently infected animals were then vaccinated intramuscularly twice, in the right hind calf muscle on day 15 and on day 25 post-infection. Animals were immunized on day 15 with 20 μg and on day 25 with 10 μg of RR2 protein mixed with 100 μg CpG/guinea pig and 150μg alum. Animals were mock-immunized with the CpG oligonucleotide (5’- TCGTCGTTGTCGTTTTGTCGTT-3’) (TriLink Inc., Santa Fe Springs, CA) using 100 μg CpG/guinea pig and 150μg alum (Alhydrogel, Accurate Chemical and Scientific Corp., Westbury, NY).

One week later, on day 32, one group was treated with an adenovirus containing CXCL11 (1x10^10^ GC) while the other group remained untreated. Mock-vaccinated guinea pigs that had received Alum + CpG adjuvants alone were used as negative control (*Mock*). From day 32 to day 42 post-infection, vaccinated and non-vaccinated guinea pigs were observed daily for (*i*) the severity of genital herpetic lesions scored on a scale of 0 to 4, and pictures of genital areas were taken; and (*ii*) vaginal swabs were collected daily from day 32 through day 42 post-infection to detect virus shedding and to quantify HSV-2 DNA copy numbers.

### Monitoring of primary or recurrent HSV-2 disease in guinea pigs

Guinea pigs were examined for vaginal lesions and were recorded for each individual animal daily on a scale of 0 to 4, where 0 reflects no disease, 1 reflects redness, 2 reflects a single lesion, 3 reflects coalesced lesions, and 4 reflects ulcerated lesions.

### qPCR for quantification of HSV-2 in Vaginal Swabs and dorsal root ganglia

Vaginal swabs were collected daily using a Dacron swab (type 1; Spectrum Laboratories, Los Angeles, CA) starting from day 35 until day 42 post-challenge. Individual swabs were transferred to a 2mL sterile cryogenic vial containing 1ml culture medium and stored at -80°C until use. On day 65 post-challenge, twelve lower lumbar and sacral dorsal root ganglia (DRG) per guinea pig were collected by cutting through the lumbar end of the spine. DNA was isolated from the collected vaginal swab and DRG of guinea pigs by using DNeasy blood and tissue kits (Qiagen). The presence and quantification of HSV- 2 was done by real-time PCR (StepOnePlus Real-Time PCR System) using 50-100 ng vaginal swab DNA or 250 ng of DRG DNA. HSV-2 DNA copy number was determined using purified HSV-2 DNA (Advanced Biotechnologies, Columbia, MD) and based on a standard curve of the *C_T_* values. The *C_T_* values from unknown samples were plotted on the purified HSV-2 DNA standard curves, and the number of HSV-2 DNA copies per assay was calculated. Each guinea pig sample was analyzed in duplicate. Samples with <150 copies/ml by 35 cycles were reported as negative. Primer and probe sequences for the HSV-2 Us9 were primers forward, 5′-GGCAGAAGCCTACTACTCGGAAA-3′, and reverse 5′-CCATGCGCACGAGGAAGT-3′, and probe with reporter dye 5′-FAM-CGAGGCCGCCAAC- MGBNFQ-3′ (FAM, 6-carboxyfluorescein). All reactions were performed using TaqMan gene expression master mix (Applied Biosystems) and data was collected and analyzed on StepOnePlus real-time PCR system.

### Splenocytes isolation

Guinea pigs’ spleens were harvested at 80 days post-infection. Briefly, spleens were placed in 10 ml of cold PBS with 10% fetal bovine serum (FBS) and 2X antibiotic–antimycotic (Life Technologies, Carlsbad, CA) and subsequently minced finely and sequentially passed through a 100 µm mesh and a 70 µm mesh (BD Biosciences, San Jose, CA). Cells were then pelleted via centrifugation at 400 × *g* for 10 minutes at 4°C. Red blood cells were lysed using a lysis buffer (ammonium chloride) and washed again. Isolated splenocytes were diluted to 1 × 10^6^ viable cells per ml in RPMI media with 10% (v/v) FBS and 2 × antibiotic–antimycotic. Viability was determined by trypan blue staining.

### Isolation of lymphocytes from guinea pigs’ vaginal mucosa

Vaginal mucosa was obtained from the guinea pigs. The genital tract was minced into fine pieces whereby the tissue pieces were then transferred into a new tube with fresh RPMI-10 containing collagenase and digested at 37 °C for two hours on a rocker set to vigorously stir the tissue. The digested tissue suspension was then passed through a 100 μm cell strainer on ice, followed by centrifugation. Lymphocytes in the cell pellets were separated using Percoll gradients by centrifugation at 900 x g, at room temperature, for 20 minutes with the brake-off. The lymphocytes at the interface layer between 40% and 70% Percoll layers were harvested, washed with RPMI 1:3, and spun down at 740 x g.

### Isolation of lymphocytes from guinea pigs’ dorsal root ganglia

Dorsal root ganglia (DRGs) were removed from the guinea pigs and minced into fine pieces whereby the tissue pieces were then transferred into a new tube with fresh RPMI-10 containing collagenase and digested at 37 °C for 1 hour on a rocker set to vigorously stir the tissue. The digested tissue suspension was then passed through a 100 μm cell strainer on ice, followed by centrifugation. Lymphocytes in the cell pellets were separated using 25% Percoll by centrifugation at 900 x g, at room temperature, for 20 minutes with the brake off. The lymphocytes were pelleted, washed with RPMI 1:3 and spun down at 740 x g.

### Isolation of lymphocytes from guinea pigs’ dorsal root ganglia

Dorsal root ganglia (DRGs) were removed from the guinea pigs and minced into fine pieces whereby the tissue pieces and transferred into a new tube with fresh RPMI-10 containing collagenase and digested at 37 °C for 1 hour on a rocker set to vigorously stir the tissue. The digested tissue suspension was then passed through a 100 μm cell strainer on ice, followed by centrifugation. Lymphocytes in the cell pellets were separated using 25% Percoll by centrifugation at 900 x g, at room temperature, for 20 minutes with the brake-off. The lymphocytes were pelleted, washed with RPMI 1:3, and spun down at 740 x g.

### Flow cytometry analysis

Vaginal mucosa cells, DRG cells, and splenocytes were analyzed by flow cytometry. The following antibodies were used: mouse anti-guinea pig CD8 (clone MCA752F, Bio-Rad Laboratories, Hercules, CA), mouse anti-guinea pig CD4 (clone MCA749PE, Bio-Rad Laboratories), anti-mouse CRTAM (clone 11-5, Biolegend, San Diego, CA), hamster anti-mouse PD- 1 clone J43, BD Biosciences, San Jose, CA), anti-mouse/human CD44 (clone IM7, Biolegend), anti-mouse CD69 (clone H1.2F3, BD Biosciences, San Jose, CA), anti-mouse CXCR3 (clone CXCR3- 173, Biolegend) and anti-mouse CD103 (clone 2E7, Biolegend). For surface staining, mAbs against various cell markers were added to a total of 1X10^6^ cells in phosphate-buffered saline containing 1% FBS and 0.1% sodium azide (fluorescence-activated cell sorter [FACS] buffer) and left for 45 minutes at 4°C. At the end of the incubation period, the cells were washed twice with FACS buffer. A total of 100,000 events were acquired by LSRII Fortessa (Becton Dickinson, Mountain View, CA).

For intracellular staining, mAbs against various cell markers were added to a total of 1 × 10^6^ cells in 1× PBS containing 1% FBS and 0.1% sodium azide (FACS buffer) for 45 minutes at 4°C. After washing with FACS buffer, cells were permeabilized for 20 minutes on ice using the Cytofix/Cytoperm Kit (BD Biosciences) and then washed twice with Perm/Wash Buffer (BD Biosciences). Intracellular cytokine IFN-gamma mAb was then added to the cells and incubated for 45 min on ice in the dark. Cells were washed again with Perm/Wash Buffer and FACS buffer and fixed in PBS containing 2% paraformaldehyde (Sigma-Aldrich, St. Louis, MO). For each sample, 100,000 total events were acquired on BD LSR II Fortessa, followed by analysis using FlowJo software (TreeStar, Ashland, OR).

Ab capture beads (BD Biosciences) were used as individual compensation tubes for each fluorophore in the experiment. To define positive and negative populations, we used fluorescence minus one control for each fluorophore used in this study when initially developing staining protocols.

### Statistical analyses

Data for each assay were compared by Student’s *t*-test using GraphPad Prism version 5 (La Jolla, CA). Differences between the groups were identified by ANOVA and multiple comparison procedures, as we previously described (33). Data are expressed as the mean ± SD. Results were considered statistically significant at *P* < 0.05.

## RESULTS

### Therapeutic immunization of HSV-2 infected guinea pigs with vaccine candidate RR2 and treatment with adenovirus containing CXCL11 protects against recurrent genital disease: G

As illustrated in **Fig. 1A**, guinea pigs (*n* = 18) were infected intravaginally with 5 x 10^5^ pfu of HSV-2 (strain MS). Once the acute infection was resolved, latently infected animals were randomly divided into two groups (*n* = 6) and then vaccinated twice intramuscularly, on days 15 and 25 post-infection with HSV-2 protein RR2 emulsified in Alum + CpG adjuvants, as detailed in *Material and Methods*. The third group (*n* = 6) received alum plus CpG adjuvants alone and were used as negative controls. One week later, one of the two RR2 vaccinated groups was treated with an adenovirus containing CXCL11, while the other group remained untreated. Starting on day 32, the guinea pigs were observed and scored regularly for genital lesions until day 42. RR2 vaccinated animals treated with CXCL11 animals exhibited considerably lower cumulative vaginal lesions than RR2 alone vaccinated but significantly lower cumulative vaginal lesions than mock vaccinated controls (*P* < 0.005, **Fig. 1B**). Similarly, a significant reduction in cumulative virus vaginal shedding was observed in animals treated with RR2/CXCL11 compared to the RR2 alone and mock-vaccinated controls (*P* < 0.005, **Fig. 1C**).

**Figure 1:**
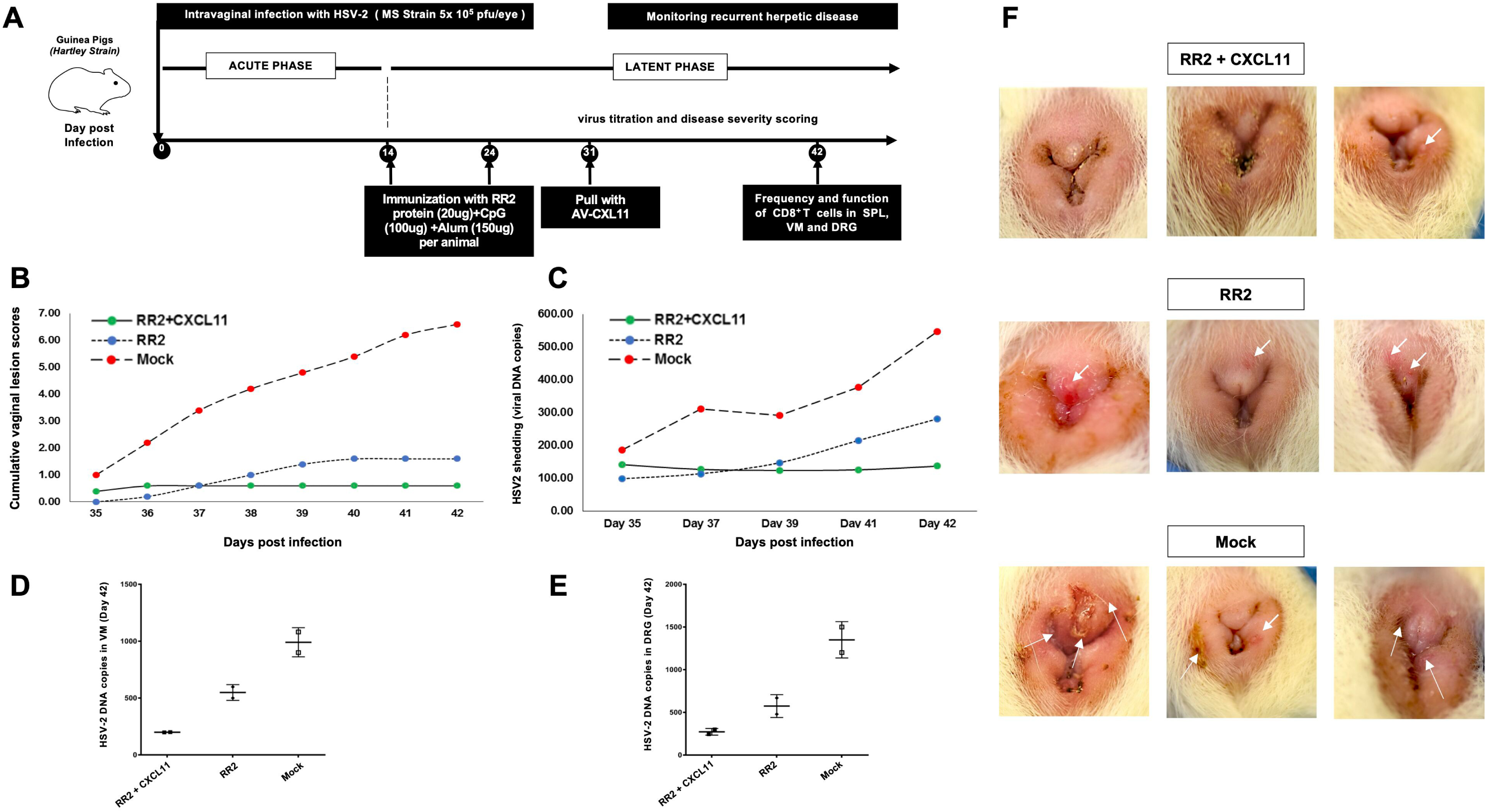
Protection against recurrent genital herpes infection and disease in HSV-2 infected guinea pigs following therapeutic prime/pull vaccination with the RR2 protein/CXCL11: (A) Timeline showing HSV-2 infection, immunization, immunological and virological analyses. Guinea pigs (*n* = 18) were infected intravaginally with 5 x 10^5^ pfu of HSV-2 (strain MS). Once the acute infection was resolved, latently infected animals were randomly divided into 2 groups (*n* = 6) and vaccinated intramuscularly twice, on days 15 and 25 post-infection with individual HSV-2 antigen RR2 emulsified in Alum + CpG adjuvants, as detailed in *Material and Methods*. One week later, on day 32, one group was treated with an adenovirus containing CXCL11, and the other group remained untreated. Mock-vaccinated guinea pigs, which received Alum + CpG adjuvants alone, were used as negative control (*Mock*). From day 32 to day 42 post-infection, vaccinated and non- vaccinated guinea pigs were observed daily for: (*i*) the severity of genital herpetic lesions scored on a scale of 0 to 4 and pictures of genital areas taken; and (*ii*) vaginal swabs were collected daily from day 32 to day 42 post-infection to detect virus shedding and to quantify HSV-2 DNA copy numbers. (**B**) Cumulative vaginal lesion (**C**) Cumulative vaginal shedding of HSV-2 during recurrent infection (viral titers). (**D**) HSV-2 DNA copy numbers were detected in the VM of vaccinated and mock vaccinated guinea pigs on day 42. (**E**) HSV-2 DNA copy numbers detected in the DRG of vaccinated and mock vaccinated guinea pigs on day 42. The severity of genital herpetic lesions was scored on a scale of 0 to 4, where 0 reflects no disease, 1 reflects redness, 2 reflects a single lesion, 3 reflects coalesced lesions, and 4 reflects ulcerated lesions. (**F**) Representative images of genital lesions in guinea pigs vaccinated with: (*i*) RR2 and treated with CXCL11 (*top 3 pictures*); (*ii*) with *RR2 alone* (*middle 3 pictures*); and (*iii*) with mock vaccinated group (*bottom 3 pictures)*. The indicated *P* values show statistical significance between the HSV-2 vaccinated and mock-vaccinated control groups *P* < 0.05 is considered significant.

On day 42 post-infection (i.e., 20 days after the second and final immunization), guinea pig vaginal swabs were collected to quantify HSV-2 virus by qPCR. The RR2/CXCL11 treated guinea pigs exhibited lower HSV-2 DNA copy numbers in the VM and DRG than mock-vaccinated controls or RR2 vaccinated alone (**Fig. 1D** and **1E**). The severity of genital herpetic lesions scored on a scale of 0 to 4, also confirmed the reduction in disease manifestation in RR2/CXCL11 treated in comparison to RR2 alone or mock vaccinated guinea pigs (**Fig. 1F**). The lowest genital lesions were observed in guinea pigs vaccinated with RR2 protein and treated with CXCL11 (**Fig. 1F**). RR2 alone displayed moderate protection whereas, the mock vaccinated group showed no protection against genital herpes lesions (**Fig. 1F**,). Altogether, these results indicate that therapeutic immunization with RR2 protein and treatment with AV containing CXCL11 protected HSV-2-seropositive guinea pigs against recurrent genital herpes infection and disease.

### Therapeutic prime/pull vaccination of HSV-2 infected guinea pigs with RR2 protein/CXCL11 increased the frequencies of tissue-resident CD4^+^ and CD8^+^ T cells

We next investigated the frequencies of CD4^+^ and CD8^+^ T cells in the spleen (SPL), vaginal mucosa (VM), and dorsal root ganglia (DRG) of guinea pigs. Guinea pigs were infected and immunized, as detailed above (**Fig 1A**). On day 42, the vaccinated and control animals were euthanized, and the frequencies of tissue-resident CD4^+^ and CD8^+^ T cells in SPL, DRG, and VM was detected by fluorescence-activated cell sorting (FACS). The gating strategy is shown in supplementary **Fig. 1**. No significant differences in the frequencies of CD4^+^ and CD8^+^ T cells in the SPL of vaccinated guinea pigs compared to the mock-vaccinated group were observed (**Fig. 2A**). However, a significantly higher frequencies of CD4^+^ and CD8^+^ T cells were observed in the VM and DRG of guinea pigs treated with CXCL11/RR2 group, compared to the mock vaccinated group (*P* < 0.05). Also, the frequency of CD4^+^ and CD8^+^ T cells in CXCL11/RR2 group was considerably higher than RR2 alone group (**Fig. 2B** and **2C**).

**Figure 2:**
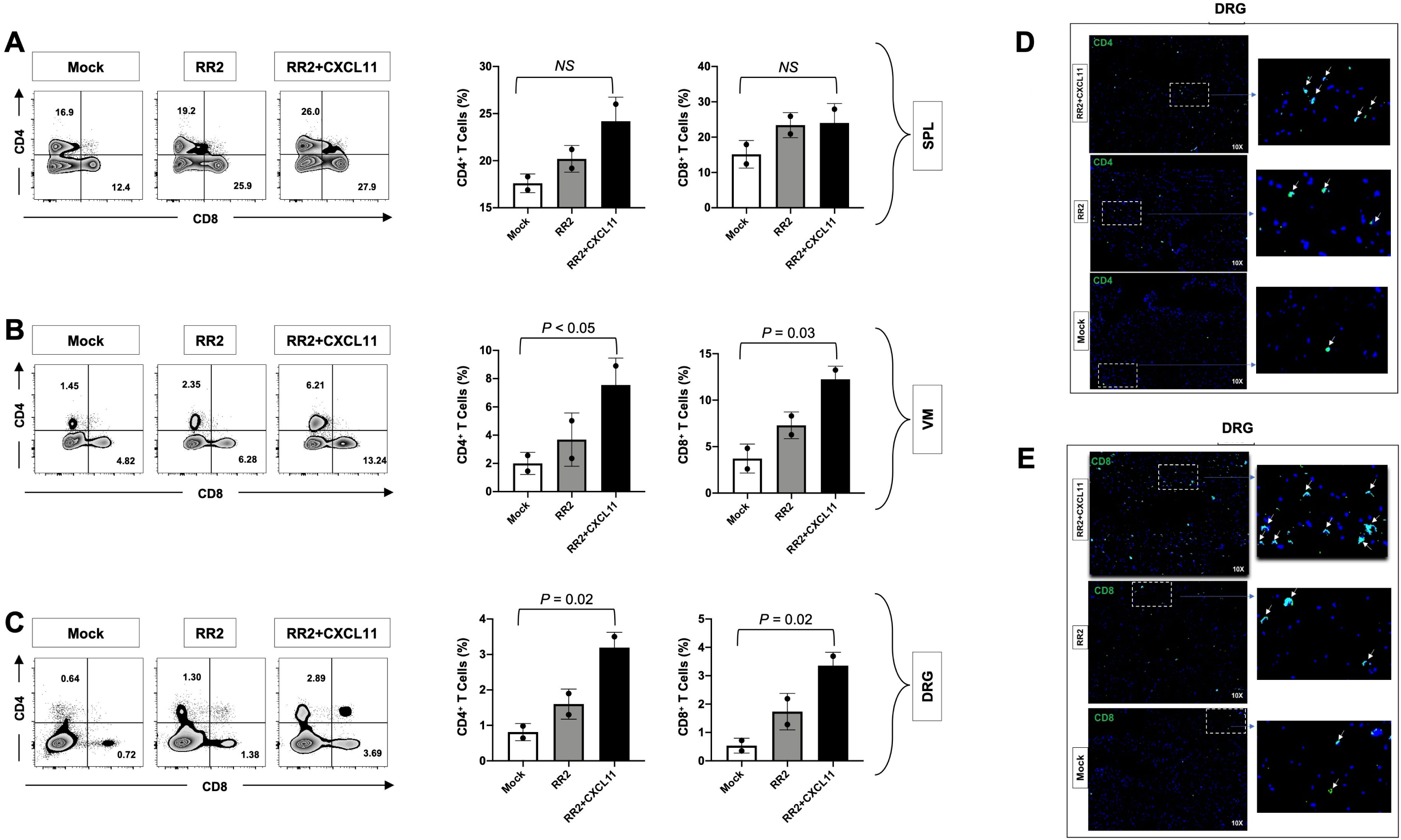
Frequencies of CD4^+^ and CD8^+^ T cells in the vaginal-mucosa and DRG of HSV-2 infected in guinea pigs following therapeutic prime/pull vaccination with RR2/CXCL11: Forty- two days post-infection, guinea pigs were euthanized, and single-cell suspensions from the spleen, vaginal mucosa, and DRG were obtained after collagenase treatment. The SPL, VM, and DRG cells were stained for CD4^+^ and CD8^+^ T cells and then analyzed by FACS. Representative FACS data (left panel) and average frequencies (right panel) of CD4^+^ and CD8^+^ T cells detected in the (**A**) SPL, (**B**) VM, and (**C**) DRG of RR2 and RR2/CXCL11 vaccinated and mock vaccinated animals. Cells were analyzed using a BD LSR Fortessa Flow Cytometry system with a total of 4 x 10^5^ events. The Zebra plot showing the percentage of CD4^+^ T cells and CD8^+^ T cells is representative of two independent experiments. The indicated *P* values performed by t-test for significance show statistical significance between prime/pull vaccinated and mock-vaccinated control groups. *P* < 0.05 is considered statistically significant. (**D**) Visualization of CD4^+^ and (**E**) CD8^+^ T cell infiltration were shown, using fluorescence microscopy, within the DRG of HSV-2 infected guinea pigs following therapeutic prime/pull vaccination with RR2 protein/CXCL11. Sections of the DRG from vaccinated and mock- vaccinated animals were co-stained using mAb specific to CD4^+^ T or CD8^+^ T cells (*Green*) and DAPI (DNA stain, *Blue*).

The frequency of CD4^+^ and CD8^+^ T cells in the DRG samples was also analyzed using fluorescence microscopy. We observed a higher CD4^+^ T cell infiltration in the DRG of CXCL11/RR2 treated guinea pigs in comparison to and RR2 alone and mock vaccinated group (**Fig. 2D**). Similarly, a higher frequency of CD8^+^ T cell infiltration was observed in the DRG of CXCL11/RR2 treated guinea pigs compared to RR2 alone and mock vaccinated group (**Fig. 2E**). RR2 alone group exhibited higher frequency of CD4 and CD8-T cells over mock vaccinated group.

### Induced protection from HSV-2 infection following therapeutic prime/pull vaccination with RR2 and treatment with adenovirus containing CXCL11 is associated with more functional tissue-resident IFN-γ^+^CRTAM^+^CD8^+^ T cells

We sought to determine the function of CD8^+^ T cells in the SPL, VM, and DRG of HSV-2 infected guinea pigs following prime/pull vaccination with RR2/CXCL11. On day 42 after the second and final therapeutic immunization and treatment with CXCL11, guinea pigs were euthanized, single-cell suspensions from the SPL, VM, and DRG tissue were obtained, and the function of SPL, VM-resident, and DRG-resident CD8^+^ T cells were analyzed for the production of IFN- γ and CRTAM expression by FACS. We found a significant difference in the frequencies of IFN- γ producing CD8^+^ T cells in the SPL of guinea pigs that were vaccinated with RR2+CXCL11 and mock vaccinated groups (**Fig. 3A**). Significantly high frequencies of IFN- γ producing CD8^+^ T cells were observed to be induced by RR2 vaccinated group treated with CXCL11, followed by the RR2 alone vaccinated group compared to those with the mock vaccinated group (i.e., adjuvant alone) in the VM and DRG tissues (*P* < 0.05) (**Fig. 3B** and **3C**).

**Figure 3.**
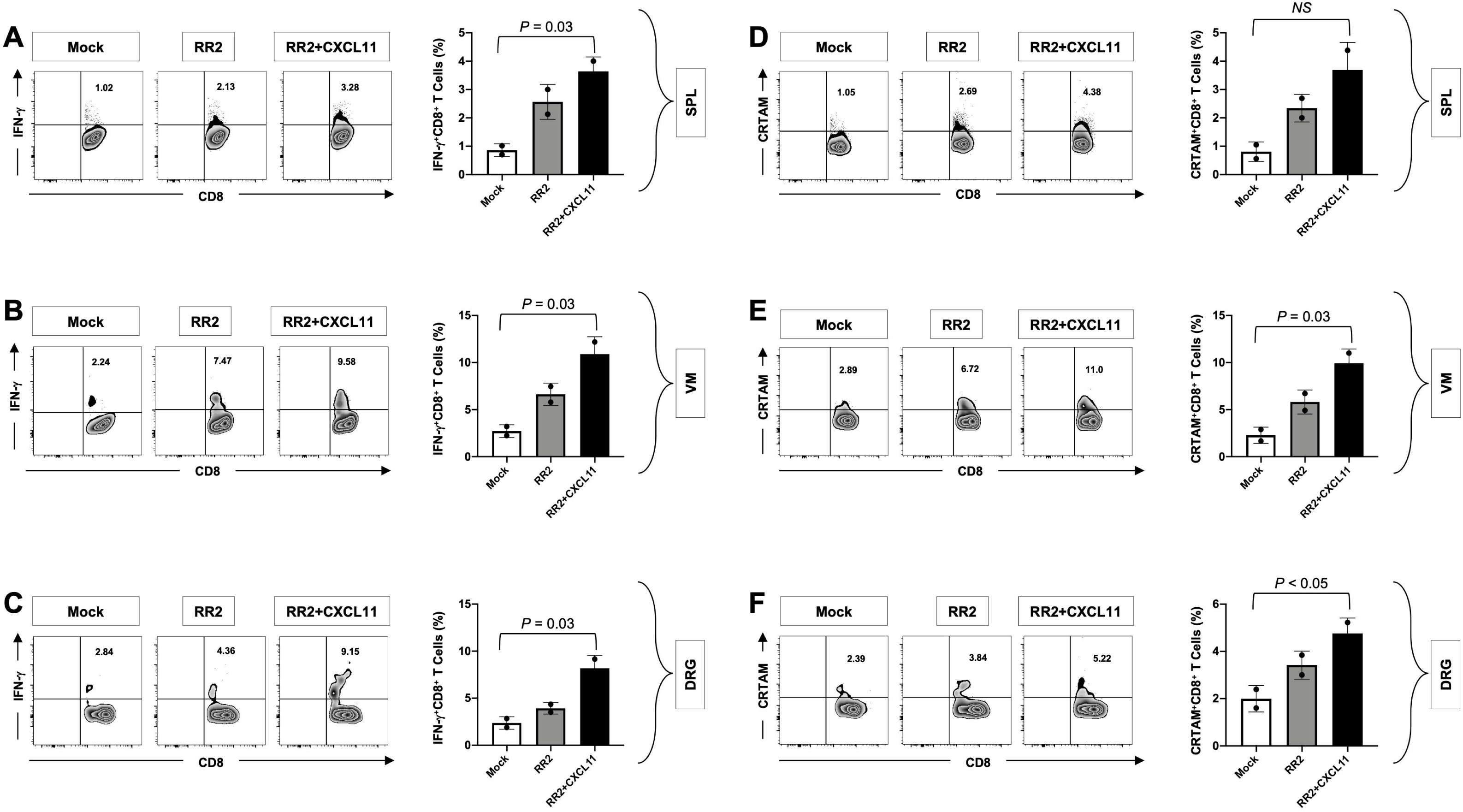
The function of tissue-resident CD8^+^ T cells from HSV-2 infected guinea pigs following therapeutic prime/pull vaccination with the RR2 protein/CXCL11: Functional analysis of CD8^+^ T cells from HSV-2-infected guinea pigs following therapeutic prime/pull vaccination with RR2 protein/CXCL11, RR2 alone and mock vaccinated animals. **Single-cell** suspension from SPL, VM and DRG were obtained post-infection and immunization. Single cells from SPL, VM, and DRG were stained for functional marker IFN- γ, activation marker CRTAM and CD8 and analyzed by FACS. Representative FACS data (left panel) and average frequencies (right panel) of IFN-γ^+^ CD8^+^ T cells detected in the (**A**) SPL, (**B**) VM, and (**C**) DRG of RR2^+^CXCL11^+^ and RR2 alone vaccinated groups and in the mock vaccinated group. The numbers on the top of each zebra plot indicate the percentage of IFN- γ^+^CD8^+^ T cells per tissue-specific total cells. Representative FACS data (left panel) and average frequencies (right panel) of CRTAM^+^ CD8^+^ T cells detected in the (**D**) SPL, (**E**) VM, and (**F**) DRG of RR2^+^CXCL11^+^ and RR2 alone vaccinated groups and in the mock vaccinated group. The numbers on the top of each zebra plot indicate the percentage of CRTAM^+^CD8^+^ T cells per tissue- specific total cells, as determined by FACS. The indicated *P* values show statistical significance between the RR2^+^CXCL11^+^ vaccinated, RR2 alone vaccinated, and mock-vaccinated control groups. *P* < 0.05 is considered significant.

Class I-Restricted T-cell-associated molecule (CRTAM) is expressed on the surface of activated CD4+ and CD8+ T-cells, by and large on the CD8^+^ T cells. It is essential for the residency of CD8+ T cells by acting as an adhesion molecule and allowing homing of T-cells in the mucosal and intraepithelial region. There was a significantly high frequency of CRTAM on CD8^+^ T cells induced by the RR2 vaccinated group treated with CXCL11, followed by vaccination by RR2 alone compared to those with the mock vaccinated group (i.e., adjuvant alone) in the VM and DRG tissues (*P* < 0.05) (**Fig. 3E** and **3F**). Conversely, we observed a non-significant but considerable difference in the frequencies of CRTAM expression on CD8^+^ T cells in the SPL of guinea pigs vaccinated with RR2/CXCL11 and RR2 alone protein compared to the mock vaccinated group (**Fig. 3D**). In conclusion, these results indicate that the prime/pull vaccination of HSV-2-infected guinea pigs using RR2/CXCL11 induced more IFN-γ^+^ CRTAM^+^ CD8^+^ T cells within the vaginal mucocutaneous and DRG tissues that is associated with significant protection against recurrent genital herpes.

### Therapeutic prime/pull RR2/CXCL11 vaccination of HSV-2 infected guinea pigs provided protection by bolstering effector and CXCR3^+^CD8^+^ T cell responses

We sought to determine the CXCR3 expression on the functional effector CD8^+^T cells. On day 42 after the final therapeutic prime/pull vaccination and treatment with CXCL11, guinea pigs were euthanized, and single-cell suspensions from the SPL, VM, and DRG tissue were obtained, and the effector function of SPL, VM- resident, and DRG-resident CD8^+^ T cells was studied by analyzing the expression of CXCR3 and CD44 on CD8+T-cells by FACS analysis. CXCR3 is a receptor for CXCL11 and is required for the mobilization of T-cells to the site of infection. This leads to attenuation in clinical manifestation of herpetic lesions in case of HSV-2 infection.

A significant difference was observed in the frequencies of CXCR3^+^CD8^+^ T cells in the SPL of guinea pigs vaccinated with RR2/CXCL11 compared to the mock vaccinated group (**Fig. 4A**). Similarly, higher frequencies of CXCR3^+^CD8^+^ T cells were observed in the RR2/CXCL11vaccinated group treated with CXCL11, followed by vaccination with the RR2 alone compared to those with the mock vaccinated group (i.e., adjuvant alone) in the VM and DRG tissues (*P* < 0.05) (**Fig. 4B** and **4C**). No significant differences were observed in the frequencies of CD44 expression on CD8^+^ T cells in the SPL and DRG of guinea pigs vaccinated with RR2/CXCL11 and RR2 protein compared to the mock vaccinated group (**Fig. 4D** and **4F**). However, significantly higher frequencies of CD44 on CD8^+^ T cells were induced by the RR2/CXCL11 vaccinated group, followed by vaccinated by RR2 alone compared to those with the mock vaccinated group (i.e., adjuvant alone) in the VM tissues (*P* < 0.05) (**Fig. 4E**). These results indicate that therapeutic vaccination of HSV-2-infected guinea pigs with the RR2 protein treated with CXCL11 attracted more CXCR3^+^ CD44^+^ CD8^+^ T cells within the vaginal mucocutaneous tissue and DRG tissue associated with significant protection against recurrent genital herpes.

**Figure 4.**
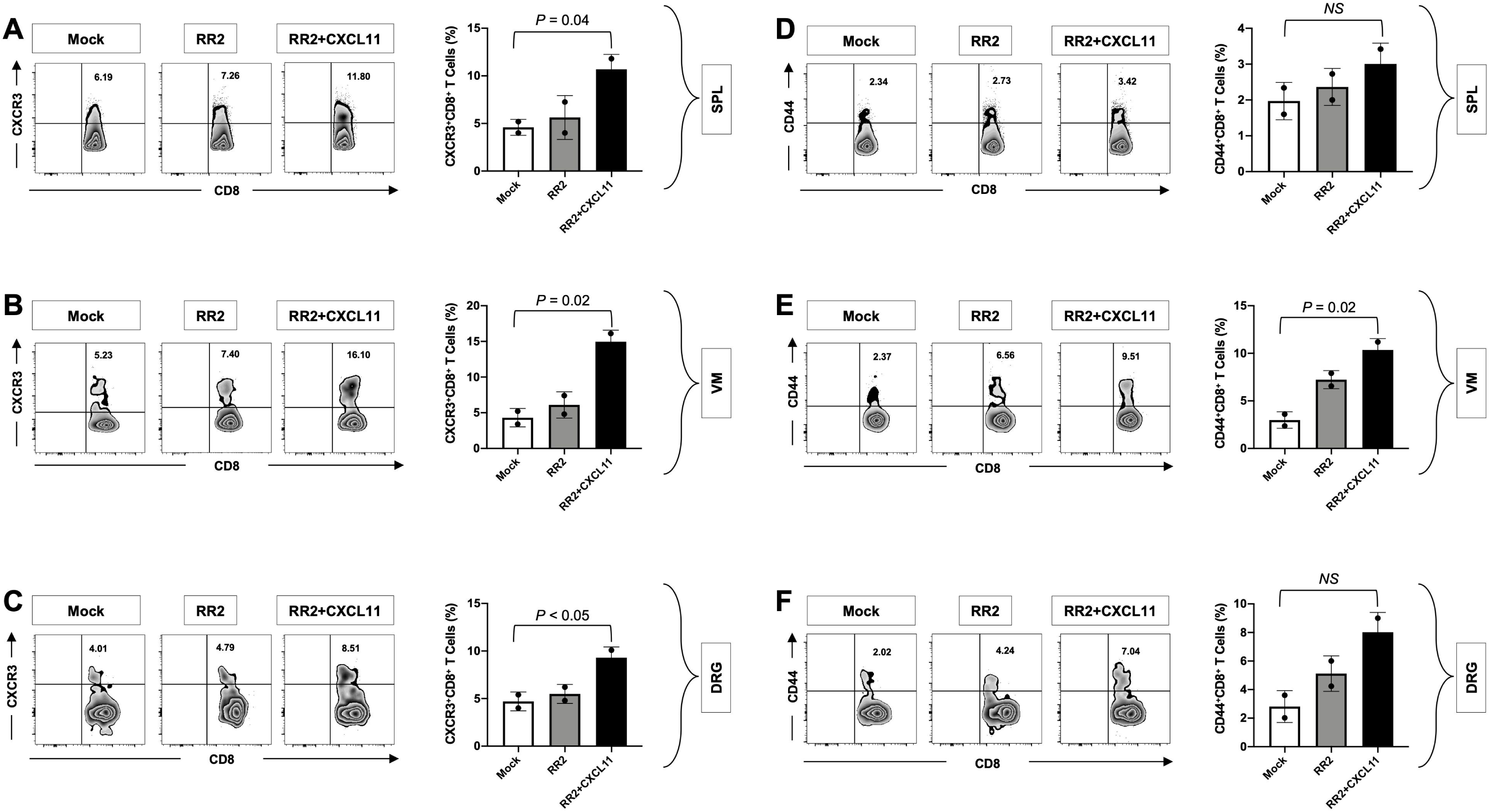
Therapeutic prime/pull vaccination of HSV-2 infected guinea pigs with RR2 protein followed by CXCL11 treatment protects by bolstering effector and CXCR3 responses: The FACS analysis is shown for CXCR3 and CD44 on CD8^+^ T cells from HSV-2-infected guinea pigs following therapeutic prime/pull vaccination with the RR2 protein/CXCL11, RR2 alone, and mock vaccinated animals. **Single-cell** suspensions from SPL, VM, and DRG were obtained post-infection and immunization. Cells were stained for CXCR3, CD44, and their CD8^+^ T cells frequencies were compared. Representative FACS data (left panel) and average frequencies (right panel) of CXCR3^+^ CD8^+^ T cells detected in the (**A**) SPL, (**B**) VM, and (**C**) DRG of RR2+CXCL11 and RR2 alone vaccinated groups and in the mock vaccinated group. The numbers on the top of each zebra plot indicate the percentage of CXCR3^+^CD8^+^ T cells per tissue-specific total cells, as determined by FACS. Representative FACS data (left panel) and average frequencies (right panel) of CD44^+^ CD8^+^ T cells detected in the (**D**) SPL, (**E**) VM, and (**F**) DRG of RR2+CXCL11 and RR2 alone vaccinated groups and in the mock vaccinated group. The numbers on the top of each zebra plot indicate the percentage of CD44^+^CD8^+^ T cells per tissue-specific total cells. The indicated *P* values show statistical significance between RR2+CXCL11 vaccinated, RR2 alone vaccinated, and mock- vaccinated control group. *P* < 0.05 is considered significant.

### Therapeutic prime/pull RR2/CXCL11 vaccination of HSV-2 infected guinea pigs efficiently generate CXCR3-dependent tissue-resident memory (T_RM_) in the VM and DRG

Tissue-resident memory or non-circulating memory cells are of prime importance against viruses as their localization in tissues gives them an enhanced capacity to mount an immediate response against viruses compared to circulating memory T-cells. For this, we sought to determine the association of various protection parameters (i.e., virus shedding and severity and frequency of recurrent genital herpes lesions) with tissue-resident memory that resides at the vaginal mucocutaneous and DRG tissue of HSV-2-infected RR2+CXCL11 and RR2 vaccinated guinea pigs.

On day 42, after the second and final therapeutic immunization and treatment with CXCL11, guinea pigs were euthanized, single-cell suspensions from the SPL, VM, and DRG tissue were obtained, and the effector function of SPL-resident, VM-resident, and DRG-resident CD8^+^ T cells was analyzed using both productions of CD69 and CD103 expression by FACS. There was no significant difference observed in the frequencies of CD69^+^CD8^+^ T cells in the SPL and DRG of guinea pigs that were vaccinated with RR2 and treated with CXCL11 and RR2 protein alone compared to the mock vaccinated group (**Fig. 5A** and **5C**). However, significantly higher frequencies of CD69^+^CD8^+^ T cells were induced by RR2 vaccinated group treated with CXCL11, followed by vaccination by RR2 alone compared to those with the mock vaccinated group (i.e., adjuvant alone) in the VM tissues (*P* < 0.05) (**Fig. 5B**).

**Figure 5.**
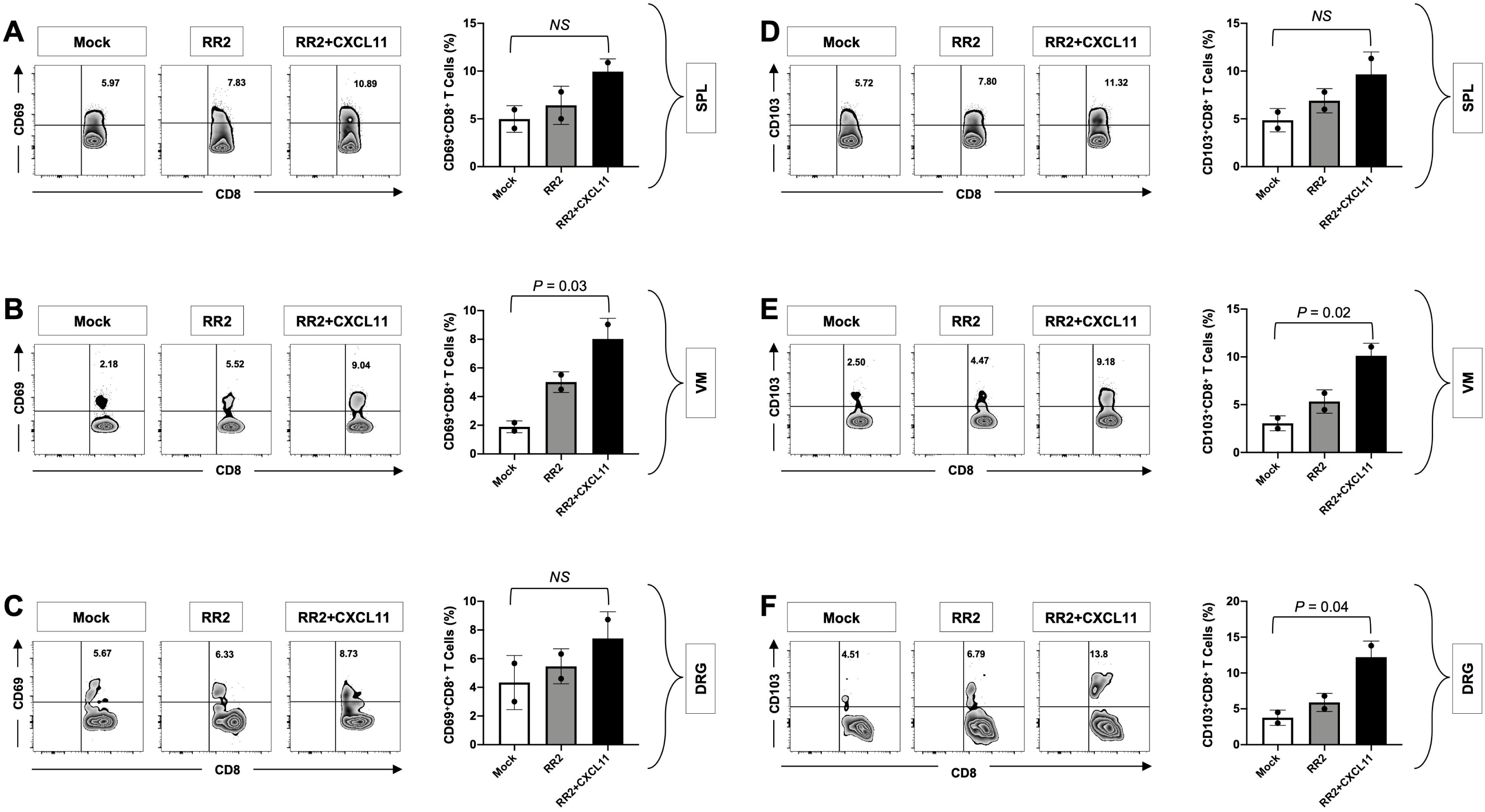
Therapeutic prime/pull vaccination of HSV-2 infected guinea pigs with RR2 *protein followed by CXCL11 treatment* efficiently generate CXCR3-dependent tissue-resident memory CD103^+^CD8^+^ T_RM_ cells in the VM and DRG: Single cell suspensions obtained post-infection and immunization from SPL, VM, and DRG were stained for memory markers CD69, CD103, and CD8 and analyzed by FACS. Representative FACS data (left panel) and average frequencies (right panel) of CD69^+^ CD8^+^ T cells were detected in the (**A**) SPL, (**B**) VM, and (**C**) DRG of RR2+CXCL11 and RR2 alone vaccinated groups and in the mock vaccinated group. The numbers on the top of each zebra plot indicate the percentage of CD69^+^CD8^+^ T cells per tissue-specific total cells, as determined by FACS. Representative FACS data (left panel) and average frequencies (right panel) of CD103^+^ CD8^+^ T cells detected in the (**D**) SPL, (**E**) VM, and (**F**) DRG of RR2+CXCL11 and RR2 alone vaccinated groups and in the mock vaccinated group. The numbers on the top of each zebra plot indicate the percentage of CD103^+^CD8^+^ T cells per tissue-specific total cells, as determined by FACS. The indicated *P* values show statistical significance between RR2+CXCL11 vaccinated, RR2 alone vaccinated, and mock-vaccinated control group. *P* < 0.05 is considered significant.

Similarly, there was no significant difference observed in the frequencies of CD103 expression gated on CD8^+^ T cells in the SPL and DRG of guinea pigs that were vaccinated with RR2/CXCL11 and RR2 protein compared to the mock vaccinated group (**Fig. 5D)**. However, significantly high frequencies of CD103^+^CD8^+^ T cells were induced by RR2 vaccinated group treated with CXCL11, followed by vaccination by the RR2 alone compared to those with the mock vaccinated group (i.e., adjuvant alone) in the VM and DRG tissues (*P* < 0.05) (**Fig. 5E** and **5F**). Taken together, these results indicate that therapeutic vaccination of HSV-2-infected guinea pigs with RR2 protein treated with CXCL11 induced higher numbers of CD8^+^ T_RM_ cells within the vaginal mucocutaneous tissue and DRG tissue associated with significant protection against recurrent genital herpes.

### Persistence of activated CD8^+^ T cells following RR2/CXCL11 vaccination of HSV-2 infected guinea pigs

During chronic viral infection, the persistence of antigens leads to impaired effector T-cell functions or T-cell exhaustion. We compared the exhaustion of CD8^+^ T cells in the SPL, VM, and DRG of HSV-2 infected guinea pigs following therapeutic prime/pull vaccination with RR2/CXCL11 and RR2 protein alone. On day 42, after the second and final therapeutic immunization and treatment with CXCL11, guinea pigs were euthanized, single-cell suspensions from the SPL, VM, and DRG tissue were obtained, and the exhaustion of SPL, VM-resident, and DRG-resident CD8^+^ T cells was analyzed using PD-1 expression by FACS. No significant differences were observed in the frequencies of PD-1^+^CD8^+^ T cells in the SPL of guinea pigs vaccinated with RR2+CXCL11 and RR2 protein compared to the mock vaccinated group (**Fig. 6A**). Similarly, there was no significant difference found in the frequencies of PD-1^+^CD8^+^ T cells by RR2 vaccinated group treated with CXCL11, followed by vaccination by RR2 alone compared to those with the mock vaccinated group (i.e., adjuvant alone) in the VM and DRG tissues (**Fig. 6B** and **6C**).

**Figure 6:**
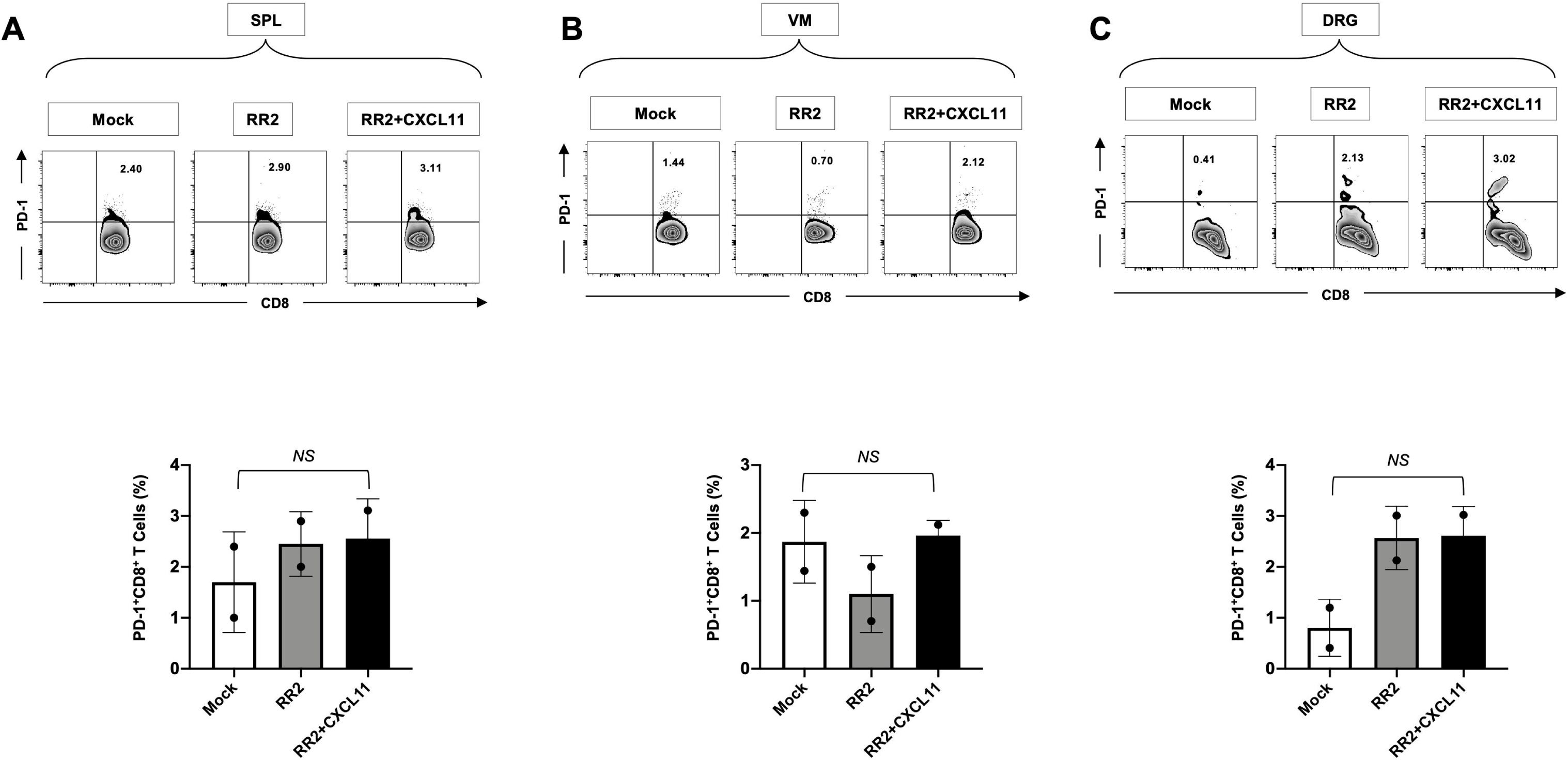
Exhaustion of CD8^+^ T cells following prime/pull therapeutic vaccination of *HSV-2 infected guinea pigs*: Single cell suspension from SPL, VM, and DRG were obtained post- infection and immunization. Single cells from SPL, VM, and DRG were stained for exhaustion marker PD-1 and CD8 and analyzed by FACS. Representative FACS data (top panel) and average frequencies (bottom panel) of PD-1^+^ CD8^+^ T cells detected in the (**A**) SPL, (**B**) VM, and (**C**) DRG of RR2+CXCL11 and RR2 alone vaccinated groups and in the mock vaccinated group. The numbers on the top of each zebra plot indicate the percentage of PD-1^+^CD8^+^ T cells per tissue-specific total cells. The indicated *P* values show statistical significance between the RR2+CXCL11 vaccinated, RR2 alone vaccinated, and mock-vaccinated control group. *P* < 0.05. is considered significant.

Overall, these results indicate that therapeutic vaccination of HSV-2-infected guinea pigs with RR2 protein treated with CXCL11 induced more functional but no difference in the exhausted CD8^+^ T_RM_ cells within the vaginal mucocutaneous tissue and DRG tissue associated with significant protection against recurrent genital herpes.

## DISCUSSION

An effective therapeutic vaccine constitutes the best and most cost-effective approach to limiting HSV-2 genital infections in humans. In one of our previous studies, we compared the protective efficacy among eight HSV-2 proteins and demonstrated that the immunization using HSV-2 RR2 protein exhibited significant protection against recurrent genital herpes disease. The protein-induced HSV-2 specific antibody and T-cell responses correlated with reduced viral shedding and disease severity. In the present study, we demonstrate the therapeutic efficacy of the prime pull strategy using protein for immunization (Prime) and a chemokine (Pull) to increase the number of HSV-2 specific T-cells in the vaginal mucosa of guinea pigs. The localized increase in the number of T-cells was done by using adenovirus expressing guinea pig-specific mucosal chemokine CXCL11. Guinea pigs were immunized (Primed) using previously studied RR2 protein and RR2-specific T-cells were pulled in VM using CXCL11. We hypothesized that the chemokine-based pulling will increase the functional, tissue-resident, IFN-g producing T-cells in the VM. Our studies demonstrate that indeed the chemokine CXCL11 treatment was able to pull the RR2-specific CD4+ and CD8+ T-cells in the VM. We observed an increase in the number and function of tissue-resident memory CD4^+^ and CD8^+^ T cells in the vaginal mucosa of guinea pigs. The pulling of T-cells manifested in a significant reduction of virus shedding and decreased recurrent genital herpes lesions. The viral reduction was significantly higher than that observed with RR2 alone.

In the present study, we evaluated the role of RR2 alone and RR2 in combination with CXCL11 and compared the protection against HSV-2 infection. Both RR2 alone and RR2/CXCL11 reduced viral shedding and demonstrated a decrease in the frequency and severity of the recurrent herpetic disease. However, guinea pigs immunized with RR2/CXCL11 displayed significantly higher protection and reduced viral numbers than RR2 alone. Comparing the immunization using protein alone (RR2) and protein with chemokine (RR2/CXCL11), we observed the latter displayed more protection against HSV-2 infection based on vaginal lesions and viral DNA copy numbers.

Chemokines like CXCL9, CXCL10, and CXCL11 are important in the adhesion and transmigration of effector T cells. Activated T-cells are recruited to peripheral tissues through the engagement of chemokine receptors. The chemokine CXCL11 binds with the receptor CXCR3 which is expressed on the surface of activated helper and cytotoxic T-cells. The receptor is maintained on the surface of these cells through the effector and memory phase. The expression of CXCR3 on activated T-cells and its ligation with its ligand (CXCL11) is important to regulate the recruitment of antigen-specific T-cells in tissues. In this regard, we evaluated the expression of CXCR3 on CD8^+^ T- cells in the VM, DRG, and spleen of guinea pigs. We observed an increased number and percentage of CXCR3^+^CD8^+^ T-cells in the VM and DRG of guinea pigs treated with CXCL11/RR2 compared to RR2 alone. The reduced frequency of CXCR3^+^CD8^+^ T-cells in mock and RR2 alone highlights the role of CXCL11/CXCR3 in the migration of CD8^+^ T-cells in the VM and DRG of CXCL11-treated guinea pigs.

Furthermore, we observed that RR2/CXCL11 significantly boosts the numbers of functional IFN-γ^+^CRTAM^+^CFSE^+^CD8^+^ T cells that were localized to healed sites of the vaginal mucocutaneous tissues. However, the treatment didn’t alter or increase the absolute number of functional CD4+T- cells in the spleen, VM or DRG of guinea pigs. The role of CXCR3 in CD4^+^ T cell migration and recruitment has been shown in other infectious models. However, in the HSV-2 guinea pig model, we believe that other chemokines might be involved in CD4^+^ T cell migration to VM and DRG. It would be of interest to identify the chemokines that pull both activated CD4^+^ and CD4^+^ T cells in the VM and DRG of infected guinea pigs.

Tissue-resident cells refer to Memory T-cells that are non-circulating and localized in the peripheral tissues. They mount an immediate response whenever a pathogen or virus infects the peripheral tissue including the female genital tract. These cells are highly heterogeneous due to their anatomic location and micro-environment-specific inflammatory cues. CD69 and CD103 are commonly used as a marker of tissue residence. Earlier studies have suggested that the application of spermicide agents to the female genital tract increased the number of CD103+Tcells and was correlated with protection. Therefore, the molecules that cause a localized increase in CD103+Cd69+T-cells may be correlated with protection. The use of RR2/CXCl11 led to the increase in CD69+CD103+T-cells in the vaginal mucocutaneous tissue.

The morbidity, socioeconomic and economic burden associated with recurrent genital herpes underscore the need for developing a therapeutic herpes vaccine (2, 34). Despite its toll, herpes has generally been seen as a “marginal” disease. Only, a handful of pharmaceutical companies and academic institutions have invested in herpes vaccine research over the past few decades due to the failure of many vaccines designed to protect from genital herpes, thus leading to disappointments in developing novel herpes vaccine strategies (35–37). In the present study, lessons learned from the failure of past clinical trials using the glycoproteins gB/gD based vaccines led us to hypothesize that a therapeutic subunit vaccine incorporating non-gB/gD “asymptomatic” HSV-2 proteins that selectively induce CD4^+^ and CD8^+^ T_RM_ cells from naturally “protected” asymptomatic women would halt or at least decrease the frequency and severity of the recurrent herpetic disease.

A reliable small animal model is ultimately the key factor for pre-clinical testing of the efficacy of the growing numbers of therapeutic herpes vaccine candidates against spontaneous herpes reactivation and recurrent disease, yet its relevance has been often overlooked. The use of genital herpes mouse models to test therapeutic vaccine candidates is precluded by the lack of spontaneous reactivation and shedding of reactivated virus in the genital tract and hence the absence of recurrent genital herpetic disease. In contrast, spontaneous reactivation occurs in HSV-2 infected guinea pigs which develop clinical and pathological features of recurrent genital herpes similar to that observed in human infections. In the guinea pig model of genital herpes, the appearance and evolution of genital lesions, the histologic changes in the genital epithelium and nerve tissues, and the establishment of latent virus infection in the DRG with the potential for reactivation, all mimic human disease. Since the phenotype and function of human blood-derived CD4^+^ and CD8^+^ T cells may not always reflect those of tissue-resident CD4^+^ and CD8^+^ T cells; the guinea pig represents the gold standard model to determine the contribution of the vaginal mucosa-tissues-resident CD4^+^ and CD8^+^ T cells versus blood-derived CD4^+^ and CD8^+^ T cells in protection against recurrent genital herpes (103). The guinea pig model has allowed us to speed up testing whether therapeutic vaccine candidates, encompassing asymptomatic protein Ags (e.g., RR2 and UL49) decrease the level of latency in DRG, spontaneous reactivation from sensory neurons of DRG, virus shedding in the genital tract and recurrent genital herpetic disease. Similar to the dl5-29 vaccine, we found the RR2 subunit vaccine to significantly boost the numbers of functional IFN-γ^+^CRTAM^+^CFSE^+^CD4^+^ and IFN-γ^+^CRTAM^+^CFSE^+^CD8^+^ T cells that were localized to healed sites of the vaginal mucocutaneous tissues associated with a significant decrease in virus shedding and a reduction in the severity and frequency of recurrent genital herpes lesions. To our knowledge, this is the first report demonstrating a correlation between, (i) a reduction in virus shedding and a decrease in recurrent genital herpes lesions induced by RR2 and, (ii) an increase in the number and function of tissue-resident memory CD4^+^ and CD8^+^ T cells in the vaginal mucosa in the guinea pig model. The phenotype memory of CD4^+^ and CD8^+^ T cells in the vaginal mucosa as to their expression of CD103, CD69, CCR7, and CD44 molecules in currently ongoing and the results will be reported in a future report.

Stress, following immunosuppression, exposure to a variety of physical or psychological/emotional stimulations, triggers HSV to reactivate from latency and virus shedding in the genital tract causes recurrent herpetic diseases (38). Asymptomatic HSV-1 and HSV-2 infections are common, pointing to the ability of the immune system to control the symptoms, despite failure to clear the virus (39). The most genital herpetic disease is due to viral reactivations from latency, rather than to primary acute infection (20). Over two-thirds of women (80%) who are seropositive for HSV-2 are asymptomatic (ASYMP) and are unaware of their infection (20). They shed and can transmit the reactivated virus but have no history of recurrent herpetic disease. In contrast, a small proportion of seropositive women are symptomatic (SYMP). They have a history of frequent and lifelong episodes of recurrent herpetic disease, often requiring doctor visitations and continuous antiviral therapy (20). It is becoming increasingly clear that the acquired herpes immunity in SYMP individuals is not sufficient for protection against virus reactivation and virus replication and disease in the genital tract (20). This implies that a successful herpes vaccine must be able to either boost the natural immune responses or initiate a protective immune mechanism that does not occur following natural exposure to the virus (15, 40, 41). Such protective immune responses might be directed against different Ags than those by the immune system of SYMP individuals. The correct response to the recent “failures” of clinical HSV vaccine trials using gB/gD glycoproteins is to encourage the testing of novel herpes tegument, capsid, and regulatory proteins as candidate vaccines.

The recurrent genital herpetic disease is one of the most common sexually transmitted diseases with a worldwide prevalence of infection predicted to be over 3.5 billion individuals for HSV-1 and over 536 million for HSV-2 (2). Moreover, HSV-1/HSV-2 recombinant strains are reported to be under a wide circulation (42). The majority of HSV-1 and HSV-2 seropositive individuals are asymptomatic and do not show any apparent recurrent symptoms, implying that many seropositive individuals can readily and quietly transmit the virus to their partners. Historically, it is harder to develop therapeutic vaccines than prophylactic vaccines because of the many immune-evasion strategies by HSV-1 and HSV-2 to evade the control by the host’s immune system in place in already infected individuals (43–45). Thus, considerable effort has been applied to developing a less-challenging prophylactic herpes vaccine to prevent herpes infection in seronegative patients. However, given the staggering numbers of already infected HSV-1 and HSV-2 individuals we contend that developing a therapeutic herpes vaccine to reduce herpes shedding and alleviate herpetic disease in symptomatic patients with recurrent outbreaks is highly desired. This global herpes prevalence stresses the urgency of developing a therapeutic vaccine.

In the present study, we rationally selected HSV proteins that were highly recognized by the immune system from ‘naturally” protected asymptomatic individuals as therapeutic vaccine candidates. The safety, immunogenicity, and protective efficacy of these eight HSV-2 proteins were assessed in a pre-clinical therapeutic vaccine trial using the guinea pig model of recurrent genital herpes. Compared to the *dl*5-29, a whole HSV-2 live vaccine that was recently tested in clinical trials (46), we found that therapeutic immunization with gD, VP22 or RR2 protein produced significant protection against recurrent genital herpes infection and disease. While RR2 has been previously reported as a major target of HSV-2-specific CD8^+^ T cells in the human (26), its antibody, CD4^+,^ and CD8^+^ T cells immunogenicity and protective efficacy against recurrent genital herpes have never been reported. Recently, a recombinant protein vaccine for herpes zoster (Shingrix) was shown to be highly effective with >90% efficacy, even in subjects >80 years of age (47, 48). Similar to the herpes simplex vaccine presented herein, the herpes zoster vaccine contains a single varicella adjuvanted protein that stimulates VZV-specific CD4^+^ T cells (and low-level CD8^+^ memory T cells) and humoral responses (49). Thus, protection can be reached against both herpes simplex (this study) and herpes zoster (47–49) by a single recombinant viral protein combined with an adjuvant with induction of strong local B and T cell responses. The present investigation shows significant improvement over the response induced by live attenuated vaccines (46).

To the best of our knowledge, this study is the first to demonstrate that intravaginal delivery of HSV-2 RR2 protein antigen: (*i*) boosted both neutralizing antibodies, that appear to cross-react with gB and gD glycoproteins, (*ii*) increase the number of functional tissue-resident IFN-γ-producing CRTAM^+^CFSE^+^CD4^+^ and CRTAM^+^CFSE^+^CD8^+^ T_RM_ cells in the vaginal mucosa; and (*iii*) protected against spontaneous recurrent genital herpes in the infected guinea pigs. The HSV-2 RR2 protein, therefore, constitutes a viable B- and T-cell candidate antigen to be incorporated in future genital herpes therapeutic mucosal vaccines. In a recent clinical trial, a candidate therapeutic HSV-2 vaccine formulation designated GEN-003, made by a mixture of a fragment of infected cell protein 4 (ICP4.2), a deletion mutant of gD2 (gD2ΔTMR), and Matrix-M2 adjuvant appeared to (*i*) boost HSV-2-specific antibody and T cell responses, as measured in the peripheral blood; and (*ii*) decrease rates of viral shedding and severity of lesions (50–52). The same level of protective immunity has yet to be reproduced in a larger phase III clinical trial.

The HSV-2 ribonucleotide reductase (RR) consists of two heterologous protein subunits. The small subunit (RR2) is a 38-kDa protein encoded by the *UL40* gene and the large subunit (RR1), designated ICP10, is a 140-kDa protein encoded by the *UL39* gene (53). The RR2 is a major target of HSV-2-specific CD8^+^ T cells in humans (26). Hensel et *al.,* recently reported that a preventive prophylactic vaccine based on a combination of UL40 and the extracellular portion of the gD (gDt) protected the non-infected guinea pig model from an intravaginal HSV2 challenge (54). It remains unclear whether the observed prophylactic protection was due to UL40 or to gD, since immunization with a single Ag which formed that combination was not compared in that study. We have now extended those studies by demonstrating that a therapeutic vaccination of HSV-2 infected guinea pigs with RR2 protein alone, boosted gB- and gD-cross-reactive neutralizing antibodies and enhanced the numbers of functional IFN- γ+CRTAM^+^CFSE^+^CD4^+^ and IFN-γ+CRTAM^+^CFSE^+^CD8^+^ T cells within the VM tissues. These robust local B- and T-cell responses were associated with a significant reduction in both virus-shedding and in the severity and frequency of recurrent genital herpes lesions. Interestingly, therapeutic immunization with the RR1 protein or in combination of the RR1 and RR2 proteins did not result in a more robust protection compared to RR2 alone (*data not shown*). These pre-clinical findings suggest the HSV-2 RR2 protein alone (without combination with gB, gD or RR1 antigens) as a viable candidate to be incorporated in the next genital herpes therapeutic mucosal vaccine to be clinically tested. Compared to the intramuscular route, the intravaginal route proved to be far better in boosting tissue-resident HSV-specific CD4^+^ and CD8^+^ T cells detected at the genital epithelium. However, the mechanisms that lead to superiority of intravaginal and intranasal routes remain to be elucidated. Nevertheless, it is surprising that the protective efficacy of the RR2 protein-based subunit vaccine delivered intravaginally in HSV-2 infected guinea pigs was comparable to that induced with the *dl*5-29, a whole HSV-2 live vaccine that was recently tested in a phase I clinical trial (46). The *dl*5-29 live vaccine completes only part of the herpes lytic cycle, without spreading to other cells, while expressing many HSV-2 Ags (46).

Historically, developing therapeutic vaccines has been more difficult than developing prophylactic vaccines, which are primarily focused on eliciting virus neutralizing antibodies before viral particles infect the vaginal epithelial cells (55). Acyclovir reduces/suppresses recurrent symptomatic disease, asymptomatic shedding, and lateral transmission to some extent, however, does not clear the infection or stop recurrent disease (34, 55). This emphasizes the need for a therapeutic vaccine (more than a prophylactic vaccine), which can boost the immune system to decrease virus shedding and reduce or eliminate herpes pathology. At present, no FDA-approved therapeutic vaccines are available. Therapeutic vaccines require induction of both antibodies and potent local CD4^+^ and CD8^+^ T cell responses at the sites of infections (i.e., vaginal mucocutaneous tissue and DRG) as demonstrated in this study. Developing T cell-based therapeutic vaccines is often confronted with multiple immune evasion strategies successfully developed by herpes viruses, the immune escape artists, to persist in their hosts’ (56–59). Among the immune evasion, mechanisms are the exhaustion of antiviral memory and the effector CD4^+^ and CD8^+^ T cells (45, 60, 61). Total or partial loss of T cell function (dysfunction) occurs following repetitive HSV latent/reactivation cycles. When T cell dysfunction develops under conditions of repetitive exposure to antigens, it is called exhaustion (as reviewed in (62)) and is usually linked with the expression of T cell co-inhibitory receptors: PD-1, TIM- 3, LAG3 (CD223), TIGIT, PSGL-1, 2B4 (CD244), GITR, CTLA-4 (CD152), CD160, and BTLA (CD272). In humans, tissue-resident CD8^+^ T cells express PD-1 and appear to be dysfunctional and simultaneously express high levels of 2 to 3 immune checkpoint receptors (e.g., PD-1, TIM-3, and LAG-3). To the best of our knowledge, this is the first report demonstrating the differential expression of exhaustion markers PD-1 and TIM-3 on guinea pig memory CD4^+^ and CD8^+^ T cells associated with a lack of protective immunity against recurrent genital herpes. The mechanisms that lead to an increased expression of PD-1 and TIM-3 on vaginal mucosa-resident CD4^+^ and CD8^+^ T cells remains to be elucidated. It is likely that repetitive/sporadic virus reactivations in non-protected animals lead to persistent antigenic stimulations of T cells at the vaginal mucosa causing the observed phenotypic and functional exhaustion of tissue-resident antiviral CD4^+^ and CD8^+^ T cells. Nevertheless, our results suggest a single or combined immune checkpoint blockade using anti-PD-1 and/or anti-TIM-3 mAbs would reverse the exhaustion of antiviral tissue-resident CD4^+^ and CD8^+^ T cells at the vaginal mucosa, and hence restore their protective function. The prospect of a novel RR1 and RR2-based therapeutic herpes vaccine combined with the immune checkpoints blockade would likely increase the efficacy and longevity of protection against recurrent herpes. Therapeutic subunit vaccination, using three of the moderately protective Ags tested in this study (i.e., VP16, VP11/12, and VP13/14) combined with an immune checkpoints blockade (i.e., PD-1 and/or TIM-3) is currently being tested using the guinea pig model in our laboratory.

Considering the wealth of data addressing the antibody responses to herpes infections/immunizations in the guinea pig model, it is surprising that there are only a few reports characterizing the phenotype, function, and exhaustion of CD4^+^ and CD8^+^ T cells in this model. One limitation is the unavailability of monoclonal antibodies (mAbs) specific to guinea pigs immune cell surface CD, cytokines, chemokines, and receptors. Compared to mice, relatively few immunological reagents are available to characterize CD4^+^ and CD8^+^ T cells in guinea pigs. Herein, we describe novel immunological reagents and approaches that quantitatively and qualitatively help characterize the phenotype, function, and exhaustion of CD4^+^ and CD8^+^ T cell responses both systemically and as they occur in the female reproductive tract of the infected and vaccinated guinea pigs. IFN-γ-ELISpot, CFSE-based proliferation, surface markers of T cell activation (CRTAM) and T cell exhaustion (PD-1 and TIM-3), intracellular cytokines. Moreover, quantitative PCR (qPCR) techniques to detect and quantify recurrent HSV-2 shedding have been optimized in the guinea pig model. Besides studying infection in genital herpes infection and disease, these immunological reagents and novel approaches are useful to study infection, immunity, and immune pathogenesis of many infectious and allergic diseases in the guinea pig model (63–65).

In summary, our study demonstrates that immunizing guinea pigs with an immunogenic HSV-2 protein RR2, followed by treatment with adenovirus expressing chemokine CXCL11 provides better protection against recurrent genital Herpes. Importantly, it seems that the strategy prevents the migration of HSV-2 from mucosa to neurons. This leads to decreased reactivation and viral shedding. We believe the CD8^+^ T-cells either reduce the neuronal infection or viral replication within neurons. In addition, the presence and increase in activated CD8^+^ T-cells in genital mucosa also suggest that viral establishment and replication may be inhibited at the site of entry. However, the exact mechanism in which the T-cells function or control the infection is yet to be determined. Furthermore, the above studies were performed 7 weeks after infection establishing the long-term immunity provided by RR2 protein and the efficacy of CXCL11 in localizing the memory cells at the site of infection. The presence and magnitude of appropriate chemokines at the site of infection are important to achieve maximum protection. This could be achieved by optimizing prime-pull for 100% efficacy. This strategy of immunotherapy if optimized well can help in recruiting and establishing resident T-cells that can provide protection not only against genital herpes but also other types of sexually transmitted diseases.

## ACKNOWLEDGEMENTS

We thank Dr. Sita Awasthi from the Infectious Disease Division, Department of Medicine, Perelman School of Medicine, University of Pennsylvania, Philadelphia, Pennsylvania, for her help with the initial experiments of this project including assessing the genital herpes infection, disease, and the neutralization antibody assays in the guinea pig model.

## REFERENCES

1. Chentoufi AA, Dhanushkodi NR, Srivastava R, Prakash S, Coulon PA, Zayou L, Vahed H, Chentoufi HA, Hormi-Carver KK, BenMohamed L. 2022. Combinatorial Herpes Simplex Vaccine Strategies: From Bedside to Bench and Back. Front Immunol 13:849515.

2. Samandary S, Kridane-Miledi H, Sandoval JS, Choudhury Z, Langa-Vives F, Spencer D, Chentoufi AA, Lemonnier FA, BenMohamed L. 2014. Associations of HLA-A, HLA-B and HLA-C alleles frequency with prevalence of herpes simplex virus infections and diseases across global populations: implication for the development of an universal CD8+ T-cell epitope-based vaccine. Hum Immunol 75:715–29.

3. Zhang X, Chentoufi AA, Dasgupta G, Nesburn AB, Wu M, Zhu X, Carpenter D, Wechsler SL, You S, BenMohamed L. 2009. A genital tract peptide epitope vaccine targeting TLR-2 efficiently induces local and systemic CD8+ T cells and protects against herpes simplex virus type 2 challenge. Mucosal Immunol 2:129–43.

4. Srivastava R, Hernandez-Ruiz M, Khan AA, Fouladi MA, Kim GJ, Ly VT, Yamada T, Lam C, Sarain SAB, Boldbaatar U, Zlotnik A, Bahraoui E, BenMohamed L. 2018. CXCL17 Chemokine-Dependent Mobilization of CXCR8(+)CD8(+) Effector Memory and Tissue- Resident Memory T Cells in the Vaginal Mucosa Is Associated with Protection against Genital Herpes. J Immunol doi:10.4049/jimmunol.1701474.

5. Zhang X, Dervillez X, Chentoufi AA, Badakhshan T, Bettahi I, BenMohamed L. 2012. Targeting the genital tract mucosa with a lipopeptide/recombinant adenovirus prime/boost vaccine induces potent and long-lasting CD8+ T cell immunity against herpes: importance of MyD88. J Immunol 189:4496–509.

6. Chentoufi AA, BenMohamed L, Van De Perre P, Ashkar AA. 2012. Immunity to ocular and genital herpes simplex viruses infections. Clin Dev Immunol 2012:732546.

7. Chentoufi AA, Benmohamed L. 2012. Mucosal herpes immunity and immunopathology to ocular and genital herpes simplex virus infections. Clin Dev Immunol 2012:149135.

8. Bettahi I, Zhang X, Afifi RE, BenMohamed L. 2006. Protective immunity to genital herpes simplex virus type 1 and type 2 provided by self-adjuvanting lipopeptides that drive dendritic cell maturation and elicit a polarized Th1 immune response. Viral Immunol 19:220–36.

9. Dasgupta G, Nesburn AB, Wechsler SL, BenMohamed L. 2010. Developing an asymptomatic mucosal herpes vaccine: the present and the future. Future Microbiol 5:1–4.

10. Zhang X, Castelli FA, Zhu X, Wu M, Maillere B, BenMohamed L. 2008. Gender-dependent HLA-DR-restricted epitopes identified from herpes simplex virus type 1 glycoprotein D. Clin Vaccine Immunol 15:1436–49.

11. 11. Dasgupta G, Chentoufi AA, Kalantari-Dehaghi M, Falatoonzadeh P, Chun S, H. LC, Felgner PL, H. HD, BenMohamed L. 2012. Immunodominant “Asymptomatic” Herpes Simplex Virus Type 1 and Type 2 Protein Antigens Identified by Probing Whole ORFome Microarrays By Serum Antibodies From Seropositive Asymptomatic Versus Symptomatic Individuals. J Virology In press.

12. 12. Chentoufi AA, Kritzer E, D. YM, Nesburn AB, BenMohamed L. 2012. Towards a Rational Design of an Asymptomatic Clinical Herpes Vaccine: The Old, The New, and The Unknown… Clinical and Developmental Immunology In Press.

13. Zhang X, Chentoufi AA, Dasgupta G, Nesburn AB, Wu M, Zhu X, Carpenter D, Wechsler SL, You S, BenMohamed L. 2009. A genital tract peptide epitope vaccine targeting TLR-2 efficiently induces local and systemic CD8+ T cells and protects against herpes simplex virus type 2 challenge. Mucosal Immunol 2:129–143.

14. Srivastava R, Coulon PG, Roy S, Chilukuri S, Garg S, BenMohamed L. 2018. Phenotypic and Functional Signatures of Herpes Simplex Virus-Specific Effector Memory CD73(+)CD45RA(high)CCR7(low)CD8(+) TEMRA and CD73(+)CD45RA(low)CCR7(low)CD8(+) TEM Cells Are Associated with Asymptomatic Ocular Herpes. J Immunol 201:2315–2330.

15. Khan AA, Srivastava R, Vahed H, Roy S, Walia SS, Kim GJ, Fouladi MA, Yamada T, Ly VT, Lam C, Lou A, Nguyen V, Boldbaatar U, Geertsema R, Fraser NW, BenMohamed L. 2018. Human Asymptomatic Epitope Peptide/CXCL10-Based Prime/Pull Vaccine Induces Herpes Simplex Virus-Specific Gamma Interferon-Positive CD107(+) CD8(+) T Cells That Infiltrate the Corneas and Trigeminal Ganglia of Humanized HLA Transgenic Rabbits and Protect against Ocular Herpes Challenge. J Virol 92.

16. Khan AA, Srivastava R, Spencer D, Garg S, Fremgen D, Vahed H, Lopes PP, Pham TT, Hewett C, Kuang J, Ong N, Huang L, Scarfone VM, Nesburn AB, Wechsler SL, BenMohamed L. 2015. Phenotypic and functional characterization of herpes simplex virus glycoprotein B epitope-specific effector and memory CD8+ T cells from symptomatic and asymptomatic individuals with ocular herpes. J Virol 89:3776–92.

17. Dervillez X, Gottimukkala C, Kabbara KW, Nguyen C, Badakhshan T, Kim SM, Nesburn AB, Wechsler SL, BenMohamed L. 2012. Future of an ‘Asymptomatic’ T-cell Epitope-Based Therapeutic Herpes Simplex Vaccine. Future Virol 7:371–378.

18. Dropulic LK, Cohen JI. 2012. The challenge of developing a herpes simplex virus 2 vaccine. Expert Rev Vaccines 11:1429–40.

19. Stanberry LR. 2004. Clinical trials of prophylactic and therapeutic herpes simplex virus vaccines. Herpes 11 Suppl 3:161A–169A.

20. Belshe PB, Leone PA, Bernstein DI, Wald A, Levin MJ, Stapleton JT, Gorfinkel I, Morrow RLA, Ewell MG, Stokes-Riner A, Dubin G, Heineman TC, Schulte JM, Deal CD. 2012. Efficacy Results of a Trial of a Herpes Simplex Vaccine. N Engl J Med 366:34–43.

21. Skoberne M, Cardin R, Lee A, Kazimirova A, Zielinski V, Garvie D, Lundberg A, Larson S, Bravo FJ, Bernstein DI, Flechtner JB, Long D. 2013. An adjuvanted herpes simplex virus 2 subunit vaccine elicits a T cell response in mice and is an effective therapeutic vaccine in Guinea pigs. J Virol 87:3930–42.

22. Johnston C, Koelle DM, Wald A. 2011. HSV-2: in pursuit of a vaccine. J Clin Invest 121:4600–9.

23. Laing KJ, Magaret AS, Mueller DE, Zhao L, Johnston C, De Rosa SC, Koelle DM, Wald A, Corey L. 2010. Diversity in CD8(+) T cell function and epitope breadth among persons with genital herpes. J Clin Immunol 30:703–22.

24. Kalantari-Dehaghi M, Chun S, Chentoufi AA, Pablo J, Liang L, Dasgupta G, Molina DM, Jasinskas A, Nakajima-Sasaki R, Felgner J, Hermanson G, BenMohamed L, Felgner PL, Davies DH. 2012. Discovery of potential diagnostic and vaccine antigens in herpes simplex virus 1 and 2 by proteome-wide antibody profiling. J Virol 86:4328–39.

25. Dasgupta G, Chentoufi AA, Kalantari M, Falatoonzadeh P, Chun S, Lim CH, Felgner PL, Davies DH, BenMohamed L. 2012. Immunodominant ‘asymptomatic’ herpes simplex virus 1 and 2 protein antigens identified by probing whole-ORFome microarrays with serum antibodies from seropositive asymptomatic versus symptomatic individuals. J Virol 86:4358–69.

26. Long D, Skoberne M, Gierahn TM, Larson S, Price JA, Clemens V, Baccari AE, Cohane KP, Garvie D, Siber GR, Flechtner JB. 2014. Identification of novel virus-specific antigens by CD4(+) and CD8(+) T cells from asymptomatic HSV-2 seropositive and seronegative donors. Virology 464–465:296-311.

27. Peng T, Zhu J, Phasouk K, Koelle DM, Wald A, Corey L. 2012. An effector phenotype of CD8+ T cells at the junction epithelium during clinical quiescence of herpes simplex virus 2 infection. J Virol 86:10587–96.

28. Zhu J, Koelle DM, Cao J, Vazquez J, Huang ML, Hladik F, Wald A, Corey L. 2007. Virus-specific CD8+ T cells accumulate near sensory nerve endings in genital skin during subclinical HSV-2 reactivation. J Exp Med 204:595–603.

29. Zhu J, Peng T, Johnston C, Phasouk K, Kask AS, Klock A, Jin L, Diem K, Koelle DM, Wald A, Robins H, Corey L. 2013. Immune surveillance by CD8alphaalpha+ skin-resident T cells in human herpes virus infection. Nature 497:494–7.

30. Awasthi S, Huang J, Shaw C, Friedman HM. 2014. Blocking HSV-2 glycoprotein E immune evasion as an approach to enhance efficacy of a trivalent subunit antigen vaccine for genital herpes. J Virol doi:JVI.01130-14 10.1128/JVI.01130-14.

31. Awasthi S, Balliet JW, Flynn JA, Lubinski JM, Shaw CE, DiStefano DJ, Cai M, Brown M, Smith JF, Kowalski R, Swoyer R, Galli J, Copeland V, Rios S, Davidson RC, Salnikova M, Kingsley S, Bryan J, Casimiro DR, Friedman HM. 2014. Protection provided by a herpes simplex virus 2 (HSV-2) glycoprotein C and D subunit antigen vaccine against genital HSV-2 infection in HSV-1-seropositive guinea pigs. J Virol 88:2000–10.

32. Awasthi S, Lubinski JM, Shaw CE, Barrett SM, Cai M, Wang F, Betts M, Kingsley S, Distefano DJ, Balliet JW, Flynn JA, Casimiro DR, Bryan JT, Friedman HM. 2011. Immunization with a vaccine combining herpes simplex virus 2 (HSV-2) glycoprotein C (gC) and gD subunits improves the protection of dorsal root ganglia in mice and reduces the frequency of recurrent vaginal shedding of HSV-2 DNA in guinea pigs compared to immunization with gD alone. J Virol 85:10472–86.

33. Khan AA, Srivastava R, Chentoufi AA, Kritzer E, Chilukuri S, Garg S, Yu DC, Vahed H, Huang L, Syed SA, Furness JN, Tran TT, Anthony NB, McLaren CE, Sidney J, Sette A, Noelle RJ, BenMohamed L. 2017. Bolstering the Number and Function of HSV-1-Specific CD8+ Effector Memory T Cells and Tissue-Resident Memory T Cells in Latently Infected Trigeminal Ganglia Reduces Recurrent Ocular Herpes Infection and Disease. J Immunol 199:186–203.

34. Kuo T, Wang C, Badakhshan T, Chilukuri S, BenMohamed L. 2014. The challenges and opportunities for the development of a T-cell epitope-based herpes simplex vaccine. Vaccine 32:6733–45.

35. Hook LM, Awasthi S, Dubin J, Flechtner J, Long D, Friedman HM. 2019. A trivalent gC2/gD2/gE2 vaccine for herpes simplex virus generates antibody responses that block immune evasion domains on gC2 better than natural infection. Vaccine 37:664–669.

36. Bernstein DI, Pullum DA, Cardin RD, Bravo FJ, Dixon DA, Kousoulas KG. 2019. The HSV-1 live attenuated VC2 vaccine provides protection against HSV-2 genital infection in the guinea pig model of genital herpes. Vaccine 37:61–68.

37. Srivastava R, Hernandez-Ruiz M, Khan AA, Fouladi MA, Kim GJ, Ly VT, Yamada T, Lam C, Sarain SAB, Boldbaatar U, Zlotnik A, Bahraoui E, BenMohamed L. 2018. CXCL17 Chemokine-Dependent Mobilization of CXCR8(+)CD8(+) Effector Memory and Tissue- Resident Memory T Cells in the Vaginal Mucosa Is Associated with Protection against Genital Herpes. J Immunol 200:2915–2926.

38. Awasthi S, Huang J, Shaw C, Friedman HM. 2014. Blocking herpes simplex virus 2 glycoprotein E immune evasion as an approach to enhance efficacy of a trivalent subunit antigen vaccine for genital herpes. J Virol 88:8421–32.

39. Lekstrom-Himes JA, Hohman P, Warren T, Wald A, Nam JM, Simonis T, Corey L, Straus SE. 1999. Association of major histocompatibility complex determinants with the development of symptomatic and asymptomatic genital herpes simplex virus type 2 infections. J Infect Dis 179:1077–85.

40. 40. Roy S, Fouladi MA, Kim GJ, Ly VT, Yamada T, Lam C, Sarain SAB, BenMohamed L. 2019. Blockade of LAG-3 Immune Checkpoint Combined with Therapeutic Vaccination Restore the Function of Tissue-Resident Anti-Viral CD8+ T Cells and Protect Against Recurrent Ocular Herpes Simplex Infection and Disease. Frontiers In Immunol In Press.

41. Lopes PP, Todorov G, Pham TT, Nesburn AB, Bahraoui E, BenMohamed L. 2018. Laser Adjuvant-Assisted Peptide Vaccine Promotes Skin Mobilization of Dendritic Cells and Enhances Protective CD8(+) TEM and TRM Cell Responses against Herpesvirus Infection and Disease. J Virol 92.

42. Koelle DM, Norberg P, Fitzgibbon MP, Russell RM, Greninger AL, Huang ML, Stensland L, Jing L, Magaret AS, Diem K, Selke S, Xie H, Celum C, Lingappa JR, Jerome KR, Wald A, Johnston C. 2017. Worldwide circulation of HSV-2 x HSV-1 recombinant strains. Sci Rep 7:44084.

43. Srivastava R, Dervillez X, Khan AA, Chentoufi AA, Chilukuri S, Shukr N, Fazli Y, Ong NN, Afifi RE, Osorio N, Geertsema R, Nesburn AB, Wechsler SL, BenMohamed L. 2016. The Herpes Simplex Virus Latency-Associated Transcript Gene Is Associated with a Broader Repertoire of Virus-Specific Exhausted CD8+ T Cells Retained within the Trigeminal Ganglia of Latently Infected HLA Transgenic Rabbits. J Virol 90:3913–28.

44. Chentoufi AA, Dervillez X, Dasgupta G, Nguyen C, Kabbara KW, Jiang X, Nesburn AB, Wechsler SL, BenMohamed L. 2012. The herpes simplex virus type 1 latency-associated transcript inhibits phenotypic and functional maturation of dendritic cells. Viral Immunol 25:204–15.

45. Chentoufi AA, Kritzer E, Tran MV, Dasgupta G, Lim CH, Yu DC, Afifi RE, Jiang X, Carpenter D, Osorio N, Hsiang C, Nesburn AB, Wechsler SL, BenMohamed L. 2011. The herpes simplex virus 1 latency-associated transcript promotes functional exhaustion of virus- specific CD8+ T cells in latently infected trigeminal ganglia: a novel immune evasion mechanism. J Virol 85:9127–38.

46. Diaz F, Gregory S, Nakashima H, Viapiano MS, Knipe DM. 2018. Intramuscular delivery of replication-defective herpes simplex virus gives antigen expression in muscle syncytia and improved protection against pathogenic HSV-2 strains. Virology 513:129–135.

47. Lal H, Cunningham AL, Heineman TC. 2015. Adjuvanted Herpes Zoster Subunit Vaccine in Older Adults. N Engl J Med 373:1576–7.

48. Lal H, Cunningham AL, Godeaux O, Chlibek R, Diez-Domingo J, Hwang SJ, Levin MJ, McElhaney JE, Poder A, Puig-Barbera J, Vesikari T, Watanabe D, Weckx L, Zahaf T, Heineman TC, Group ZOES. 2015. Efficacy of an adjuvanted herpes zoster subunit vaccine in older adults. N Engl J Med 372:2087–96.

49. Leroux-Roels I, Leroux-Roels G, Clement F, Vandepapeliere P, Vassilev V, Ledent E, Heineman TC. 2012. A phase 1/2 clinical trial evaluating safety and immunogenicity of a varicella zoster glycoprotein e subunit vaccine candidate in young and older adults. J Infect Dis 206:1280–90.

50. Van Wagoner N, Fife K, Leone PA, Bernstein DI, Warren T, Panther L, Novak RM, Beigi R, Kriesel J, Tyring S, Koltun W, Lucksinger G, Morris A, Zhang B, McNeil LK, Tasker S, Hetherington S, Wald A. 2018. Effects of Different Doses of GEN-003, a Therapeutic Vaccine for Genital Herpes Simplex Virus-2, on Viral Shedding and Lesions: Results of a Randomized Placebo-Controlled Trial. J Infect Dis 218:1890–1899.

51. Bernstein DI, Wald A, Warren T, Fife K, Tyring S, Lee P, Van Wagoner N, Magaret A, Flechtner JB, Tasker S, Chan J, Morris A, Hetherington S. 2017. Therapeutic Vaccine for Genital Herpes Simplex Virus-2 Infection: Findings From a Randomized Trial. J Infect Dis 215:856–864.

52. Flechtner JB, Long D, Larson S, Clemens V, Baccari A, Kien L, Chan J, Skoberne M, Brudner M, Hetherington S. 2016. Immune responses elicited by the GEN-003 candidate HSV-2 therapeutic vaccine in a randomized controlled dose-ranging phase 1/2a trial. Vaccine 34:5314–5320.

53. Smith CC, Peng T, Kulka M, Aurelian L. 1998. The PK domain of the large subunit of herpes simplex virus type 2 ribonucleotide reductase (ICP10) is required for immediate-early gene expression and virus growth. J Virol 72:9131–41.

54. Hensel MT, Marshall JD, Dorwart MR, Heeke DS, Rao E, Tummala P, Yu L, Cohen GH, Eisenberg RJ, Sloan DD. 2017. Prophylactic Herpes Simplex Virus 2 (HSV-2) Vaccines Adjuvanted with Stable Emulsion and Toll-Like Receptor 9 Agonist Induce a Robust HSV-2- Specific Cell-Mediated Immune Response, Protect against Symptomatic Disease, and Reduce the Latent Viral Reservoir. J Virol 91.

55. Dasgupta G, Chentoufi AA, Nesburn AB, Wechsler SL, BenMohamed L. 2009. New concepts in herpes simplex virus vaccine development: notes from the battlefield. Expert Rev Vaccines 8:1023–35.

56. Banks TA, Rouse BT. 1992. Herpesviruses--immune escape artists? Clin Infect Dis 14:933–41.

57. Sharma S, Rajasagi NK, Veiga-Parga T, Rouse BT. 2014. Herpes virus entry mediator (HVEM) modulates proliferation and activation of regulatory T cells following HSV-1 infection. Microbes Infect 16:648–60.

58. Suryawanshi A, Veiga-Parga T, Rajasagi NK, Reddy PB, Sehrawat S, Sharma S, Rouse BT. 2011. Role of IL-17 and Th17 cells in herpes simplex virus-induced corneal immunopathology. J Immunol 187:1919–30.

59. Sehrawat S, Suvas S, Sarangi PP, Suryawanshi A, Rouse BT. 2008. In vitro-generated antigen-specific CD4+ CD25+ Foxp3+ regulatory T cells control the severity of herpes simplex virus-induced ocular immunoinflammatory lesions. J Virol 82:6838–51.

60. Allen SJ, Hamrah P, Gate D, Mott KR, Mantopoulos D, Zheng L, Town T, Jones C, von Andrian UH, Freeman GJ, Sharpe AH, BenMohamed L, Ahmed R, Wechsler SL, Ghiasi H. 2011. The role of LAT in increased CD8+ T cell exhaustion in trigeminal ganglia of mice latently infected with herpes simplex virus 1. J Virol 85:4184–97.

61. Mott KR, Bresee CJ, Allen SJ, BenMohamed L, Wechsler SL, Ghiasi H. 2009. Level of herpes simplex virus type 1 latency correlates with severity of corneal scarring and exhaustion of CD8+ T cells in trigeminal ganglia of latently infected mice. J Virol 83:2246–54.

62. Wherry EJ. 2011. T cell exhaustion. Nat Immunol 12:492–9.

63. Wodal W, Schwendinger MG, Savidis-Dacho H, Crowe BA, Hohenadl C, Fritz R, Bruhl P, Portsmouth D, Karner-Pichl A, Balta D, Grillberger L, Kistner O, Barrett PN, Howard MK. 2015. Immunogenicity and protective efficacy of a Vero cell culture-derived whole-virus H7N9 vaccine in mice and guinea pigs. PLoS One 10:e0113963.

64. Swanson EC, Gillis P, Hernandez-Alvarado N, Fernandez-Alarcon C, Schmit M, Zabeli JC, Wussow F, Diamond DJ, Schleiss MR. 2015. Comparison of monovalent glycoprotein B with bivalent gB/pp65 (GP83) vaccine for congenital cytomegalovirus infection in a guinea pig model: Inclusion of GP83 reduces gB antibody response but both vaccine approaches provide equivalent protection against pup mortality. Vaccine 33:4013–8.

65. Gillis PA, Hernandez-Alvarado N, Gnanandarajah JS, Wussow F, Diamond DJ, Schleiss MR. 2014. Development of a novel, guinea pig-specific IFN-gamma ELISPOT assay and characterization of guinea pig cytomegalovirus GP83-specific cellular immune responses following immunization with a modified vaccinia virus Ankara (MVA)-vectored GP83 vaccine. Vaccine 32:3963–70.

